# On the mapping between evolutionary scenarios and governing rules in state-dependent speciation-extinction models

**DOI:** 10.64898/2026.09.24.754193

**Authors:** Albert C. Soewongsono, Michael J. Landis

## Abstract

Biologists often wish to explain why one trait is more common than its alternatives among species in a clade: is it due to asymmetries in diversification or character evolution processes? State-dependent speciation–extinction (SSE) models provide a flexible framework for jointly modeling such processes. Here, we derive mathematical results that define functional relationships (“rulesets”) among rate parameters, establish a mapping between rulesets and evolutionary scenarios, and characterize how these scenarios give rise to different long-term tip state distributions. In particular, we show that multiple evolutionary scenarios can produce identical stationary distributions of tip state frequencies. Performing inference on simulated data from extant-only trees, we find that not all scenarios are equally favored for explaining datasets constructed solely from extant species. This bias could arise from the interplay between mixing rates, elapsed time, and the conditioning inherent in tree likelihoods. As a proof of concept, we apply our framework to the binary SSE (BiSSE) model across multiple Squamate clades and conduct a literature survey of previous BiSSE analyses on Angiosperm phylogeny. We show that, to some extent, the preference for certain scenarios observed in simulations is also reflected in analyses of empirical trees. This study provides a theoretical ground to directly link data-generating process and observed tip state frequencies under an SSE framework.

## 1 Introduction

Understanding why a trait is common versus rare among closely related species is of broad interest to ecologists and biologists (MacArthur 1965; Gaston 2000; Mittelbach et al. 2007; Pontarp et al. 2019). Depending on which biological system is being studied, the character of interest might be a phenotypic trait, a geographical location, or a genotypic variant. From an evolutionary perspective, the frequency of a trait also depends on when it first originated; how many species retained, gained, or lost it; and whether the trait influences how rapidly species originate or go extinct. As such, phylogenetic approaches are routinely used to reconstruct how patterns of species richness and trait variation were generated. Phylogenetic data, which include topological relationships among species, branch lengths that represent evolutionary time, and character data among sampled taxa, hold important information to infer the most likely scenario to explain the present-day distribution of traits among species.

Beginning with the pioneering work in the 1990s by Sean Nee and co-authors (Nee et al. 1992, 1994), a myriad of phylogenetic models that study diversification (speciation and extinction processes) have been developed (Morlon et al. 2024). These models compute the likelihood of a phylogenetic tree using information such as the timing of branching events, tree topology, and, in more recent models, character state information. Then, using either the frequentist approach (e.g., maximum likelihood) or Bayesian approach (e.g., Bayesian phylogenetic), one finds parameter values or the posterior distribution for each parameter in the model given the data. When the presence or absence of character data at the tips of a phylogeny is considered, a unified model that jointly models lineage diversification and character evolution is required. One such framework is the State-dependent Speciation-Extinction (SSE) models (Maddison et al. 2007; Goldberg and Igić 2012). Over the years, researchers have built on these models. On the modeling side, this is done by adding more realism that reflects empirical process as much as possible. For example, starting with SSE model with only a single binary character (e.g., BiSSE; Maddison et al. 2007), several other SSE models have emerged, such as one for multi-state characters (e.g., MuSSE; FitzJohn 2010), for modeling biogeographical processes (GeoSSE; Goldberg et al. 2011), and for accounting for variation caused by ‘hidden’ (unobserved) traits (HiSSE; Beaulieu and O’Meara 2016), among many others.

This is to say that having a high-quality phylogenetic tree is essential for correctly reconstructing phylogenetic histories of trait-dependent diversification. On the statistical side, much has been done to improve the accuracy of rate estimation despite working with partial data (i.e., extant-only phylogeny). For example, specifying the correct sampling conditions for the tree likelihood (Stadler 2013) or by incorporating fossils (do Rosario Petrucci et al. 2025) have been shown to improve diversification rate estimation. However, trees can be very uncertain, in terms of relationships and/or divergence times. One can have issue with low support nodes and inaccurate divergence time estimation. Maximum likelihood approaches quantify uncertainty by bootstrapping on the sequence data to put a confidence interval on phylogeny (Felsenstein 1985), whereas in Bayesian inference, a joint estimation of the model parameters and posterior distribution of trees (Yang and Rannala 1997) is performed. In the extreme case, we may not be able to reconstruct a phylogenetic relationship at all, due to reasons such as insufficient sequence data (Rokas et al. 2003), hybridization (Mallet et al. 2016), and recombination (Posada and Crandall 2002) that could lead to uncertainty in tree topology or a non-tree like representation (e.g., phylogenetic network).

This study is motivated by the following question: in the absence of a phylogenetic tree, what can be inferred about the data-generating process underlying the present-day observations, while still accounting for the fact that the data arise from a diversification process on an unobserved tree? Previous studies have explored evolutionary inference in the absence of a resolved phylogenetic tree by integrating over possible tree histories under an assumed branching process (Crawford and Suchard 2013), while others have examined how phylogenetic information can complement frequency-based observations in inferring evolutionary processes (May and Rannala 2024). Here, we consider a related problem in the context of state-dependent diversification by asking what information about the underlying evolutionary process is retained in present-day state frequencies alone. For example, imagine a case where there are fewer species with a hypothetical character state *A* than species with state *B*, is this imbalance in state frequencies arose from an evolutionary scenario with positive net diversification rate (i.e., speciation rate minus extinction rate) in *A* and negative net diversification rate in *B* while having higher rate of transition from *A* to *B* or other scenario with negative net diversification in *A* and positive net diversification in *B* instead? Furthermore, if multiple scenarios that correspond to the identical species state frequencies exist, how can we distinguish between them in the absence of a tree?

To answer these questions, we start by deriving a formal mathematical relationship between an evolutionary scenario and a functional relationship between model parameters from an SSE model, which we express as a “ruleset”. Here, an evolutionary scenario summarizes key relationships among evolutionary processes, such as net diversification and transition directionality, whereas a ruleset specifies the complete system of parameter relationships compatible with a given pattern of stationary state frequencies. Later in the study, we generalize this by establishing a bijection between every evolutionary scenario under a particular SSE model with their corresponding ruleset. Furthermore, we show that each of these rulesets is distinct between each other and unique to their corresponding evolutionary scenario. As a proof-of-concept, we apply in the context of BiSSE and GeoSSE (see Supplement). However, the concept is extensible to other discrete-state SSE models.

Through mathematical derivation, we show the existence of multiple evolutionary scenarios producing the same long-run pattern of tip state frequencies. Note that, since the phylogeny is absent, *pattern* here does not refer to the topological relationships among species (e.g., species diversity patterns), but rather the long-run proportions of species in different states. Hence, this paper examines a problem distinct from those addressed in previous studies of identifiability in diversification models (Louca and Pennell 2020; Dragomir et al. 2023; Tarasov and Uyeda 2024; Truman et al. 2025). The bijection between evolutionary scenarios and rulesets allows us to systematically characterize the alternative evolutionary processes that can explain the same stationary frequencies and to compare their relative prevalence. That is, we show that, as a forward-in-time process, not all scenarios occupy equal space in the set of valid trees, even though each distinct scenario can produce identical tip state frequencies.

In particular, we found, through simulations, that most scenarios derived from assuming a supercritical process comprises the majority of valid trees. This finding can be used to guide prior choice in Bayesian phylogenetics, where we want to assign sensible prior that would produce valid trees. As backward-in-time process performed on simulated dataset, we found that not all scenarios are equally preferably from inference using trees with only extant species. Scenarios assuming supercritical process are often selected, which may be due to using extant phylogeny and conditions imposed on the likelihood. We then examine this preference further by conducting empirical analyses across families and infraorders in the reconstructed squamate phylogeny and by reviewing existing studies that applied BiSSE to angiosperm phylogenies. We found that while the results are mostly consistent between simulated and empirical studies, some scenarios are either underestimated or overestimated. We comment that this discrepancy could be caused by the interplay between the convergence rate of each process and the duration over which the process has evolved in each system. In general, we found that the ranking of scenarios is not always one-to-one when the processes are considered forward and backward in time.

## 2 Methods

Here, we first define an evolutionary scenario in terms of key relationships among evolutionary processes, namely for net diversification within states and transitions between states. We then introduce the notion of a ruleset, which specifies the mathematical relationships among rate parameters in an SSE model that are compatible with a given pattern of stationary state frequencies. We establish a bijection between rulesets and evolutionary scenarios, allowing each mathematical ruleset to be interpreted in terms of its corresponding evolutionary processes. Through simulation experiments, we assess the ability of each scenario to generate valid trees. In this case, these are non-extinct trees and trees that satisfy a predefined range of imbalance in tip state frequencies. Lastly, using both simulated and empirical datasets, we assess whether certain evolutionary scenarios are preferentially supported by the data.

Analyses using an SSE model may support a hypothesis in which species possessing one character state experience positive net diversification, whereas species possessing an alternative state experience negative net diversification, with transitions occurring more frequently toward the state experiencing net species gain. These relationships among process rates collectively define an evolutionary scenario.

### Definition 1

Given an SSE model, an evolutionary scenario is expressed in terms of each net diversification rate as being positive, negative, or equal to zero, and in terms of one or more state-transition rates as being larger, smaller, or equal one or more other state-transition rates.

For example, under a BiSSE model, *λ*_*i*_ is the speciation rate in state *i, µ*_*i*_ is the extinction rate in state *i, q*_*ij*_ is the transition rate from state *i* into *j*, and the net diversification rate is *λ*_*i*_ − *µ*_*i*_ in state *i*. Per Definition 1, the set of equations {*λ*_*A*_ − *µ*_*A*_ *>* 0, *λ*_*B*_ − *λ*_*B*_ *<* 0, *q*_*AB*_ = *q*_*BA*_} would specify a valid evolutionary scenario, whereas the set of equations {*λ*_*A*_ − *µ*_*A*_ *> q*_*BA*_, *λ*_*B*_ − *λ*_*A*_ *< µ*_*B*_ − *µ*_*A*_, *q*_*AB*_ = 0} would not.

The same evolutionary scenario can also be represented more precisely by a system of equalities and inequalities among rate parameters that is compatible with a specific set of asymptotic stationary state frequencies under an SSE model. We refer to this precise representation as a ruleset. In the following sections, we develop a mathematical framework for deriving rulesets that are compatible with a given pattern of stationary state frequencies and subsequently establish their correspondence with evolutionary scenarios.

### 2.1 Mathematical background

Given a discrete-state SSE model with some underlying stationary frequencies of species in each state (e.g., geographical ranges, phenotypic traits), we can derive the rate parameters associated with the given stationary frequencies using Lemma 4 in Soewongsono and Landis (2024), which was originally developed for the GeoSSE model (Goldberg et al. 2011). Note that “stationarity” here refers to the asymptotic composition of character state frequencies among extant lineages under the branching process, rather than to the stationary distribution of the continuous-time Markov chain governing character-state transitions along individual lineages. That is:

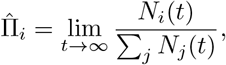

where *N*_*i*_(*t*) and *N*_*j*_(*t*) represent the number of extant lineages that are in state *i* and *j*, respectively.

In Theorem 1 we provide a generalization to Lemma 4 in Soewongsono and Landis (2024) for any discrete-state SSE model, with or without cladogenetic and anagenetic events.

#### Theorem 1

*Given a discrete-state SSE model with state space S, set of stationary frequencies*, 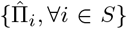, *and initial state frequencies*, Π_*i*_(0), *any set of model parameters that realizes those stationary frequencies must satisfy the following conditions:*

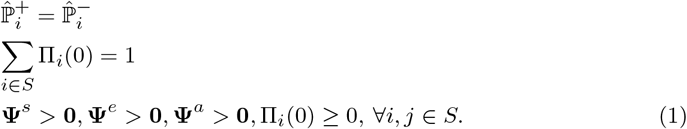

*Moreover, to classify distinct evolutionary rulesets associated with the stationary frequencies, we additionally require that, for every pair of states i, j* ∈ *S:*

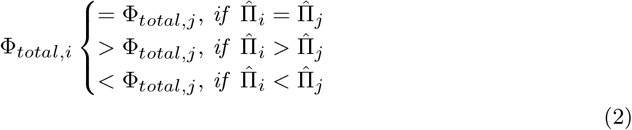

*where* 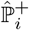 *and* 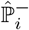 *are defined to be the probabilities of gaining or losing a new species in state i, respectively*. Φ_*total,i*_ *is defined to be the total rates of all events occurring in state i*. **Ψ**, **Ψ**^*e*^, *and* **Ψ**^*a*^ *are vectors of rates corresponding to speciation, extinction, or anagenetic change events from the model, respectively*. **0** *represents vector of zeros with corresponding size*.

**Proof:** Proof of Theorem 1 is given in Supplement 1.1. □

Solving the system of equations and inequalities in Theorem 1 will either return no solution or a ruleset(s). For the scenario without solution, it implies that there is no set of parameter values that corresponds to the particular set of stationary frequencies. We give a definition of a ruleset in Definition 2.

#### Definition 2

Given an SSE model with state space *S*, model parameters **Ψ** = {**Ψ**^*s*^, **Ψ**^*e*^, **Ψ**^*a*^}, and a set of stationary frequencies 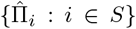, a ruleset ***ℛ*** = {*E, I*} is a collection of mathematical conditions on the model parameters that is consistent with the given stationary frequencies, where *E* and *I* denote sets of equalities and inequalities, respectively. For a given set of stationary frequencies, each ruleset, if it exists, defines a parameter solution space that is disjoint from those of all other rulesets and contains infinitely many parameter combinations.

We demonstrate Definition 2 in Section 2.1.1 for BiSSE and Section 1.4 for GeoSSE.

#### 2.1.1 Rulesets in a BiSSE model

In this section, we show how to derive a ruleset from a given set of stationary frequencies of species in different states under the BiSSE model (Maddison et al. 2007).

##### Corollary 1

*Given a BiSSE model with state space S* = {*A, B*} *and a set of stationary frequencies* 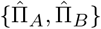 *with initial state frequencies* {*Π*_*A*_(0), Π_*B*_ (0) : Π_*A*_(0) + Π_*B*_ (0) = 1}, *we obtain a ruleset(s) by solving the following system of equations:*

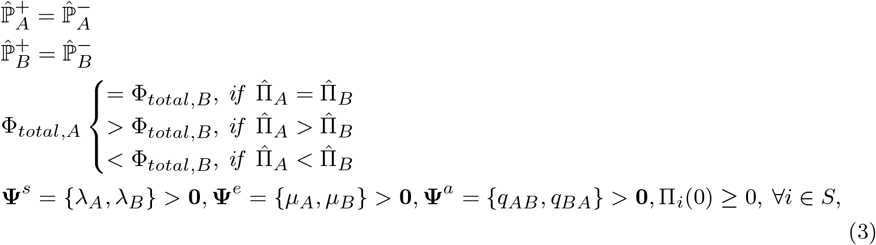

*where λ*_*A*_ *and λ*_*B*_ *are speciation rates for species in state A and B, respectively. µ*_*A*_ *and µ*_*B*_ *are extinction rates for species in state A and B, respectively. q*_*AB*_ *and q*_*BA*_ *are anagenetic rates of state changes from A to B and from B to A, respectively. For the BiSSE model, we have*

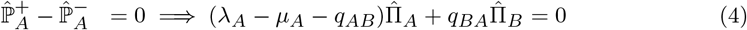

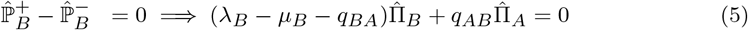

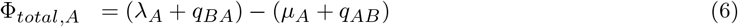

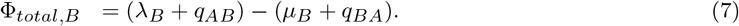

*Furthermore, we add extra constraints that net diversification rates* (*λ*_*i*_ − *µ*_*i*_) ≠0, ∀*i* ∈ *S. Otherwise, only anagenetic rates contribute to change in state frequencies over time. Note that Corollary 1 inherently excludes cases where only the directionality in transition rates drive the frequencies. That is, the method excludes (1) cases with equal net diversification in A and B* (*λ*_*A*_ − *µ*_*A*_ = *λ*_*B*_ − *µ*_*B*_) *or (2) cases where net diversification rates in A and B are both positive or negative. We refer to these cases as “trivial”, because the net diversification rates either do not differ between states or do not act in opposing directions. All other cases are referred to as “non-trivial”*.

**Proof:** Proof of Corollary 1 is given in Supplement 1.2. □

We demonstrate Corollary 1 in Example 1.

*Example 1* Suppose we have the following stationary frequencies under a BiSSE model, 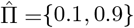. Then, applying Corollary 1, we have two rulesets associated with the frequencies 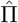, given as follows:

**Ruleset 1**

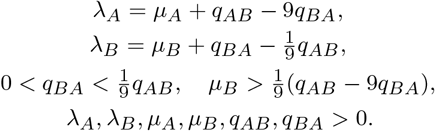

**Ruleset 2**

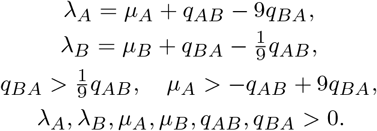

For example, the following parameters *λ*_*A*_ = 0.65, *λ*_*B*_ = 0.05, *µ*_*A*_ = 0.2, *µ*_*B*_ = 0.1, *q*_*AB*_ = 0.9, *q*_*BA*_ = 0.05 satisfy Ruleset 1 but do not satisfy Ruleset 2. On the other hand, *λ*_*A*_ = 0.1, *λ*_*B*_ = 0.3, *µ*_*A*_ = 1.0, *µ*_*B*_ = 0.2, *q*_*AB*_ = 0.9, *q*_*BA*_ = 0.2 satisfy Ruleset 2 but does not satisfy Ruleset 1. Under both cases, they correspond to the same stationary frequencies 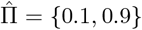 using methods given in Maddison et al. (2007) and Soewongsono and Landis (2024).

We also apply Theorem 1 to derive ruleset under GeoSSE (Supplement 1.4). Note that, by construction, Corollary 1 only includes cases in which one state is a “source” (supercritical, *λ*_*i*_ *> µ*_*i*_) and the other is a “sink” (subcritical, *λ*_*i*_ *< µ*_*i*_). When both states are sources or both are sinks, an additional condition on the overall lineage growth is required. To characterize this behavior, we introduce *ρ*, the asymptotic exponential growth rate of the total number of lineages. When *ρ >* 0, the system is supercritical, with the expected number of lineages growing asymptotically while the state frequencies converge to a stationary distribution. When *ρ <* 0, the system is subcritical, with the expected number of lineages declining asymptotically. In both cases, the relative state frequencies remain governed by the balance between diversification and anagenetic transitions, as shown in Corollary 2.

##### Corollary 2

*Given a BiSSE model with state space S* = {*A, B*}, *a set of stationary frequencies* 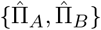 *with initial state frequencies* {*Π*_*A*_(0), Π_*B*_ (0) : Π_*A*_(0) + Π_*B*_ (0) = 1}, *and asymptotic exponential growth rate ρ of change in the total species count across all states, we obtain a ruleset(s) by solving the following system of equations:*

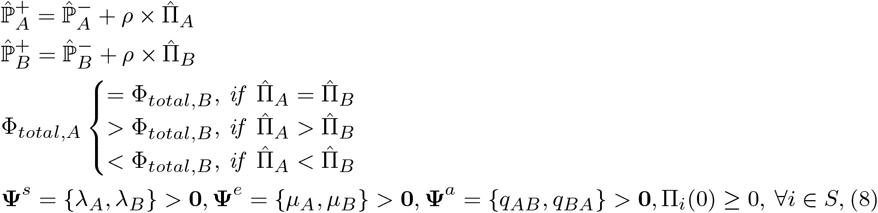

*where λ*_*A*_ *and λ*_*B*_ *are speciation rates for species in state A and B, respectively. µ*_*A*_ *and µ*_*B*_ *are extinction rates for species in state A and B, respectively. q*_*AB*_ *and q*_*BA*_ *are anagenetic rates of state changes from A to B and from B to A, respectively. For the BiSSE model, we have:*

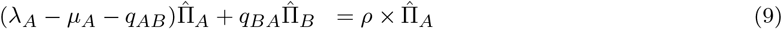

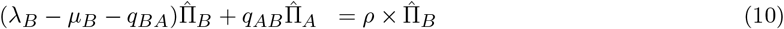

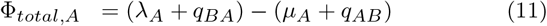

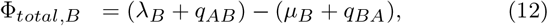

*where supercritical growth occurs when*

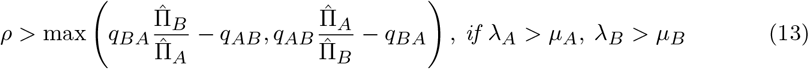

*and subcritical growth occurs when*

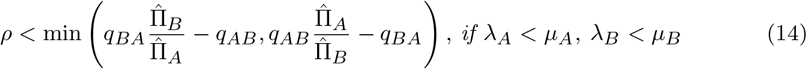

*Furthermore, we add extra constraints that net diversification rates* (*λ*_*i*_ − *µ*_*i*_) ≠ 0, ∀*i* ∈ *S. Note the left-hand side terms of* Eqs. (9)−(10) *represent the mean growth matrix, M, for a branching process, where*

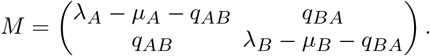

**Proof:** Proof of Corollary 2 is given in Supplement 1.3. □

It follows that Corollary 1 is a special case of Corollary 2, assumming *ρ* = 0. Note that *ρ* = 0 does not imply that the number of lineages is constant over time; rather, it implies that the expected number remains constant, while individual realizations may exhibit substantial stochastic fluctuations.

### 2.2 Symmetries between rulesets

Here we formally define symmetry between two rulesets, which correspond to two sets of stationary frequencies with permuted state labels under an SSE model. For simplicity, we present the definition in the context of the BiSSE model, although the generalization to other SSE models is straightforward. We begin by formally defining a symmetrical pair of stationary state frequencies, given in Definition 3.

#### Definition 3

Let

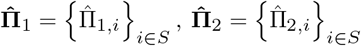

be stationary state frequencies from an SSE model defined on the state space *S*. We say 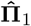 and 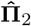 form a **symmetrical pair** if there exits a bijection

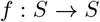

such that, for every *i* ∈ *S*,

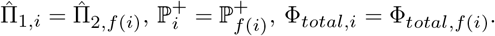

For example, under BiSSE, 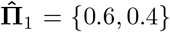 and 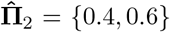 form a symmetrical pair. However, under GeoSSE, 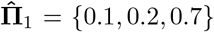 and 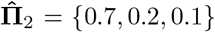 are not a symmetrical pair because 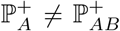 and Φ_*total,A*_ ≠ Φ_*total,AB*_ by re-labelling of indices, according to Corollary S1. Finally, we define symmetry between rulesets in\ Definition 4. Furthermore, Lemma 1 establishes a relationship between symmetry in stationary state frequencies and symmetry in rulesets.

#### Definition 4

We say two rulesets are symmetric if we can derive one ruleset from another by swapping the indices on rate parameters.

#### Lemma 1

*Given a pair of symmetric stationary state frequencies, their rulesets are also symmetric*.

**Proof:** Proof of Lemma 1 is given in Supplement 1.6 □

As an immediate consequence of Lemma 1, two rulesets are distinct if and only if they are not symmetrical. Additionally, the number of rulesets is invariant for any pair of symmetrical stationary state frequencies. That is, both 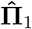 and 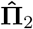 will have the same number of rulesets by consequence from Lemma 1. This consequence can be seen in Figure S1 for BiSSE and Figure S2 for a 2-region GeoSSE.

Using examples, we use Lemma 1 in the context of a BiSSE model (Example 2) and a 2-region GeoSSE model (Example S3; Supplement).

*Example 2* Suppose a BiSSE model with state space *S* = *{A, B}* with **Ψ**^*s*^ = *{λ*_*A*_, *λ*_*B*_ *} >* **0, Ψ**^*e*^ = *{µ*_*A*_, *µ*_*B*_ *} >* **0, Ψ**^*a*^ = *{q*_*AB*_, *q*_*BA*_*} >* **0**. Additionally, the net divesification rates in both states are non-zero. That is, *λ*_*A*_ − *µ*_*A*_ ≠ 0 and *λ*_*B*_ − *µ*_*B*_ ≠ 0. Then, given a pair of symmetric state frequencies as follows:

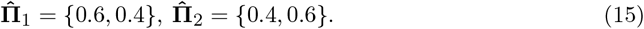

Then, using Corollary 1 (solved using `Mathematica` (Wolfram Research, Inc. 2023)), rulesets that correspond to 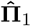 are as follows:

1. **Ruleset 1**

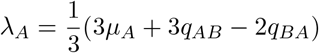

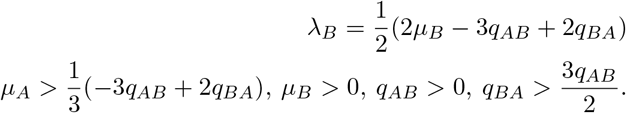
2. **Ruleset 2**

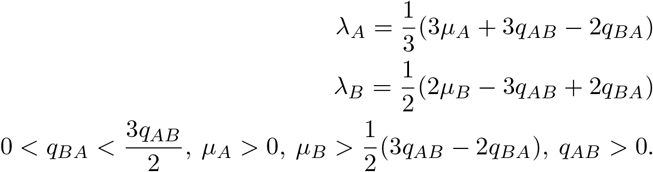

On the other hand, rulesets that correspond to 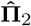 are as follows:

1. **Ruleset 1**

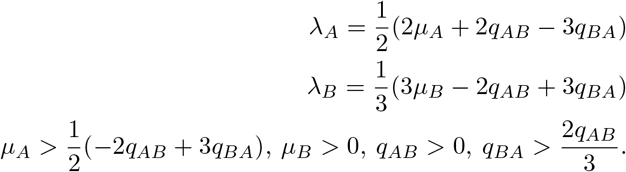
2. **Ruleset 2**

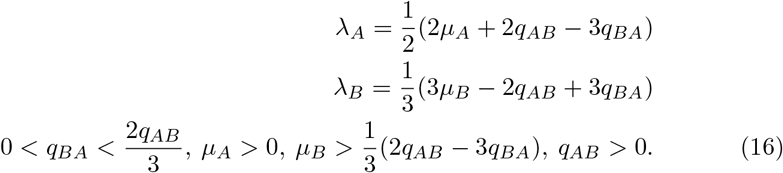

It is clear that ruleset 1 in 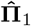 is symmetric to ruleset 2 in 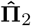 after swapping the indices on rate parameters. Similarly, ruleset 2 in 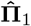 is symmetric to ruleset 1 in 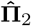. □

### 2.3 Counting distinct rulesets across different state frequencies

We quantify the number of distinct non-trivial rulesets corresponding to a given set of stationary frequencies under an SSE model, based on the approach developed in Theorem 1. We examine this result in the context of a BiSSE model using Corollary 1 (see Fig. S1) and a 2-region GeoSSE model (Corollary S1; Fig. S2).

1. We draw sets of stationary frequencies independently from a 2-dimensional meshgrid (BiSSE case) and from a 3-dimensional meshgrid (2-region GeoSSE case).
2. For each set of stationary frequencies, we find the number of distinct non-trivial rulesets that correspond to the given set using Corollary 1 (BiSSE case) and using Corollary S1 (2-region GeoSSE case).
3. We plot the ruleset counts for all sets of stationary frequencies as a ternary plot.

#### Corollary 3

*Suppose we have a BiSSE model with state space S* = *{A, B} with rate parameters λ*_*A*_, *λ*_*B*_, *µ*_*A*_, *µ*_*B*_, *q*_*AB*_, *q*_*BA*_ *>* 0. *Given a set of stationary frequencies* 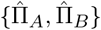 *and if either* 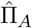 *or* 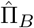 *is equal to zero, then there is no ruleset that satisfies the system of equations described in Corollary 1*.

**Proof:** Proof of Corollary 3 is given in Supplement 1.8. □

We also derive the GeoSSE analogue of Corollary 3, given as Corollary S4.

### 2.4 Rulesets as distinct evolutionary scenarios

We establish a bijective map between a ruleset and its corresponding evolutionary scenario for a given SSE model. This bijection represents a direct evolutionary interpretation for each ruleset compatible with a given stationary frequency: each ruleset corresponds to a distinct evolutionary scenario and ranges of SSE model parameters that asymptotically produce a particular set of stationary state frequencies. Thus, identical stationary frequencies do not necessarily imply identical underlying evolutionary processes. The mapping allows us to systematically characterize the alternative evolutionary processes that are consistent with a given set of stationary frequencies. As a proof-of-concept, we demonstrate this idea in the context of a BiSSE model. However, it can be readily extended to any discrete SSE model.

Recall also that we distinguish between trivial and non-trivial rulesets based on the relationships between net diversification rates. As described following Corollary 1, trivial rulesets have net diversification rates that either do not differ between states or do not act in opposing directions, whereas all other rulesets are non-trivial. Regardless of this distinction, a ruleset and its corresponding evolutionary scenario represent the same relationships at different levels of description. A ruleset specifies the mathematical relationships among model parameters that are compatible with a given directionality of stationary state frequencies (see Definition 2). In contrast, an evolutionary scenario provides a qualitative interpretation of those parameter relationships in terms of evolutionary processes, such as net lineage gain or loss within states and the favored direction of transitions between states (see Definition 1). For example, in BiSSE, given stationary state frequencies with 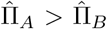, a ruleset is defined by the parameter relationships *λ*_*A*_ *> µ*_*A*_, *λ*_*B*_ *< µ*_*B*_, and *q*_*AB*_ *> q*_*BA*_, and through additional constraints on their maximum and minimum values. The corresponding evolutionary scenario interprets these relationships as net species gain in state *A*, net species loss in state *B*, and transitions being favored from *A* to *B*.

As seen from Examples 1 and S2, multiple distinct rulesets can correspond to identical stationary frequencies. We postulate that each of these rulesets corresponds to a specific evolutionary scenario. For example, the parameter set *λ*_*A*_ = 0.65, *λ*_*B*_ = 0.05, *µ*_*A*_ = 0.2, *µ*_*B*_ = 0.1, *q*_*AB*_ = 0.9, and *q*_*BA*_ = 0.05 belongs to Ruleset 1 in Example 1 and corresponds to an evolutionary scenario in which there is a net gain of species in state *A* and a net loss of species in state *B*. In contrast, the parameter set *λ*_*A*_ = 0.1, *λ*_*B*_ = 0.3, *µ*_*A*_ = 1.0, *µ*_*B*_ = 0.2, *q*_*AB*_ = 0.9, and *q*_*BA*_ = 0.2 belongs to Ruleset 2 in Example 1 and corresponds to an evolutionary scenario in which there is a net loss of species in state *A* and a net gain in state *B*.

Before we establish the mapping between rulesets and evolutionary scenarios, we first derive a general statement for obtaining the different rulesets within each directionality class of stationary state frequencies under a BiSSE model, as defined in Definition 5, and formally stated in Theorem 2.

#### Definition 5

Given a forward-in-time tree that has evolved under a BiSSE model with parameters ***θ***, the different directionalities between state frequencies (referred to as directionality classes), whether defined by the stationary frequencies 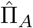 and 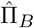 or by the observed frequencies at the tips of the tree, *f*_*A*_ and *f*_*B*_, are as follows:

1. **Directionality class a:** 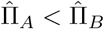 (resp. *f*_*A*_ *< f*_*B*_).
2. **Directionality class b:** 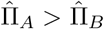 (resp. *f*_*A*_ *> f*_*B*_).
3. **Directionality class c:** 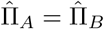 resp. *f*_*A*_ = *f*_*B*_).

#### Theorem 2

*Suppose a BiSSE model where all the rate parameters are positive with non-zero net diversification rates. Given a set of non-zero stationary frequencies*, 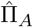 *and* 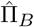, *we have two non-trivial rulesets for any pair of stationary frequencies within each directionality class for stationary frequencies, as defined in Definition 5. They are as follows:*

1. *For* 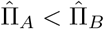, *the two non-trivial rulesets are:* ***Ruleset 1a***

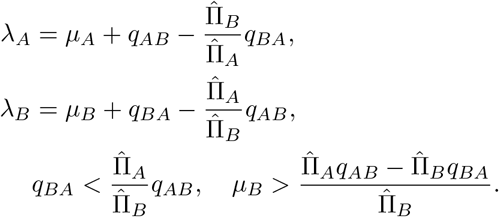

***Ruleset 2a***

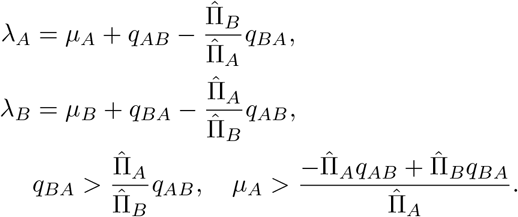
2. *For* 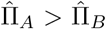, *the two non-trivial rulesets are given by the corresponding rulesets for* 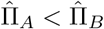 *by interchanging the state labels A and B:* ***Ruleset 1b***

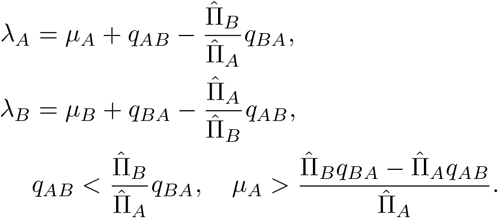

***Ruleset 2b***

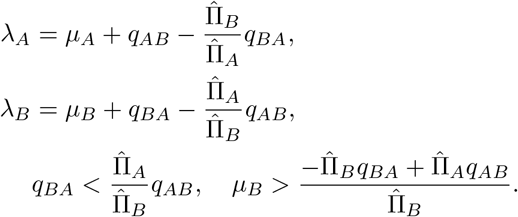
3. *For* 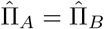, *the two rulesets are:* ***Ruleset 1c***

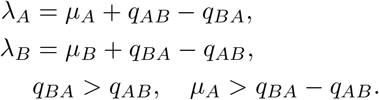

***Ruleset 2c***

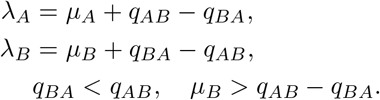

**Proof:** Proof of Theorem 2 is given in Supplement 1.10. □

Finally, using Theorem 2, we define a bijection that maps each ruleset within a directionality class to a specific evolutionary scenario, as given in Lemma 2.

#### Lemma 2

*Suppose a BiSSE model where all the rate parameters are positive with non-zero net diversification rates. Given a set of stationary frequencies*, 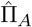 *and* 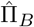, *for each directionality class defined in Definition 5, we have the following bijection g* : *ruleset i* → *non-trivial evolutionary scenario i, given as follows:*

1. 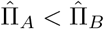 *directionality class (non-trivial):*
  - *Ruleset 1a* → *species net gain in A, species net loss in B, and more transitions into B (scenario 1a)*.
  - *Ruleset 2a* → *species net loss in A, species net gain in B (scenario 2a)*. *Rulesets 1b and 2b for the* 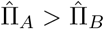 *directionality class, together with their corresponding evolutionary scenarios, are obtained from Rulesets 1a and 2a and their corresponding scenarios, respectively, by interchanging the state labels A and B*.
2. 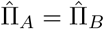 *directionality class (non-trivial):*
  - *Ruleset 1c* → *species net loss in A, species net gain in B, and more transitions into A (scenario 1c)*.
  - *Ruleset 2c* → *species net gain in A, species net loss in B, and more transitions into B (scenario 2c)*. *In addition to those non-trivial evolutionary scenarios above, we also have the following trivial evolutionary scenarios for each directionality class, which can be obtained using Corollary 2, as follows:*
3. 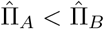 *directionality class (trivial):*
  - *Ruleset 3a* → *species net gain in A, species net gain in B, and more transitions into B (scenario 3a)*.
  - *Ruleset 4a* → *species net gain in A, species net gain in B, and more transitions into A (scenario 4a)*.
  - *Ruleset 5a* → *species net loss in A, species net loss in B, and more transitions into B (scenario 5a)*.
  - *Ruleset 6a* → *species net loss in A, species net loss in B, and more transitions into A (scenario 6a)*.
  - *Ruleset 7a* → *equal net diversification in A and B, and more transitions into B (scenario 7a)*. *Rulesets 3b–7b for the* 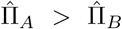 *directionality class, together with their corresponding evolutionary scenarios, are obtained from Rulesets 3a-7a and their corresponding scenarios, respectively, by interchanging the state labels A and B*.
4. 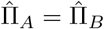 *directionality class (trivial):*
  - *Ruleset 3c* → *equal speciation rates, equal extinction rates, and equal transition rates (scenario 3c)*.

**Proof:** Proof of Lemma 2 is given in Supplement 1.11. □

### 2.5 Relaxation time to stationarity

We have showed that multiple distinct scenarios (rulesets) can induce identical stationary state frequencies. However, such scenarios still differ in how quickly they approach stationarity. The spectral gap, *γ*, of a matrix defines the rate at which system reaches convergence (i.e., effective strength toward convergence). In our case, we are interested in the spectral gap of the matrix governing the expected abundance of lineages in each state.

The inverse of the spectral gap, 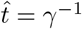, defines a relaxation timescale toward stationarity. Note that this differs from the expected waiting time required for the system to reach stationarity. Since the birth-death process is stochastic, the expected time to stationarity depends on the initial state of the system and the chosen error tolerance between the stationary and observed state frequencies. On the other hand, the relaxation time is an intrinsic property of the underlying process and does not depend on either the tolerance or the initial state. Specifically, the relaxation time characterizes the timescale over which deviations from the stationary frequencies decay, with larger values of 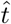 indicating slower convergence and vice versa. More generally, the spectral gap determines the asymptotic rate at which deviations from stationarity decay, rather than the absolute distance to stationarity at a particular time. For a given underlying process, this distance decreases asymptotically according to

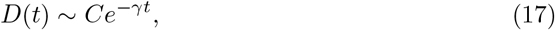

where *D*(*t*) defines the absolute distance between the observed frequencies at time *t* and the stationary frequencies under the given parameter values, and *C* is a constant that depends on the initial state and the underlying parameters of the process. Thus, larger values of *γt* generally correspond to greater convergence toward stationarity within a given process. Consequently, processes with faster convergence rates may still exhibit substantial deviations from stationarity if the tree is young, whereas processes with slower convergence rates may nonetheless appear close to stationarity if the tree is sufficiently old. However, the absolute distance at a particular value of *γt* can differ among processes because the constant *C* depends on the model parameters. Consequently, a process with a larger value of *γt* need not be closer to stationarity than a different process with a smaller value of *γt*. Nevertheless, within a given process, faster convergence (larger *γ*) or a longer elapsed time *t* leads to more relaxation toward the stationary frequencies.

From Corollary 2, we can define the relaxation time as follows:

#### Corollary 4

*Given a BiSSE model with state space S* = *{A, B}, M denotes the state-abundance matrix governing the expected lineage dynamics in each state. That is:*

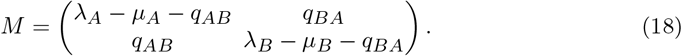

*M has a unique, real-value dominant eigenvalue ρ which defines the asymptotic growth rate of the system. The rate of convergence (spectral gap) is defined as:*

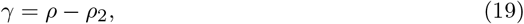

*where ρ > ρ*_2_ *such as ρ is the dominant eigenvalue and ρ*_2_ *is the second largest eigenvalue. Moreover*,

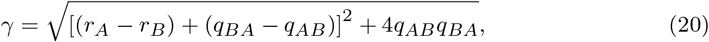

*where r*_*A*_ = *λ*_*A*_ − *µ*_*A*_ *and r*_*B*_ = *λ*_*B*_ − *µ*_*B*_. *In other words, r*_*A*_ *and r*_*B*_ *define net diversification rates in both states. By definition, the relaxation time*, 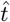, *is defined as γ*^−1^.

**Proof:** Proof of Corollary 4 is given in Supplement 1.12. □ The spectral gap as defined in Eq. (20) can be viewed as a mixture of two components. We refer to (*r*_*A*_ − *r*_*B*_) +(*q*_*BA*_ − *q*_*AB*_) as the net asymmetry term, which captures the combined asymmetry in net diversification and transition rates between states, and 4*q*_*AB*_*q*_*BA*_ as the bidirectional mixing term, which captures the contribution of transitions occurring in both directions. For example, under Scenario 4a (or 4b) in which we have competing forces between diversification pushing lineage accumulation in one direction (*r*_*A*_ *< r*_*B*_) and transitions between states pushing in the opposite direction (*q*_*BA*_ *> q*_*AB*_), the net asymmetry term will approach zero when there is a balance between these two forces (i.e., *r*_*A*_ − *r*_*B*_ ≈ *q*_*BA*_ − *q*_*AB*_), and the spectral gap under this scenario is only modulated by its mixing term. In this case,

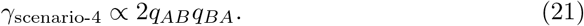

On the other hand, under Scenario 2a (or b), where diversification exhibits source-sink dynamics (*r*_*A*_ *< r*_*B*_) and transition between states can push in either direction (*q*_*BA*_ *> q*_*AB*_ or *q*_*BA*_ *< q*_*AB*_), the net asymmetry term can either reinforce or partially offset the diversification asymmetry. When transition asymmetry aligns with diversification asymmetry (*r*_*A*_ *< r*_*B*_ and *q*_*BA*_ *< q*_*AB*_), the magnitude of this term increases, leading to faster convergence (larger *γ*); when they oppose each other, it reduces the convergence rate.

### 2.6 Assessing statistical power for each evolutionary scenario

Here, we aim to assess: (1) the ability of each evolutionary scenario to generate valid trees, and (2) which evolutionary scenarios are preferentially inferred when fitting the model to extant-only trees. We address these questions in the context of a BiSSE model, but the application to other SSE models is straightforward. Moreover, to assess the robustness of our results, we perform the analysis using datasets simulated under multiple settings, as summarized in Table 1. In all simulation scenarios, we simulated 100 valid extant-only trees using slow rates and another 100 valid trees using fast rates for each tree size *{*25, 50, 100, 200, 400, 800}. We choose these tree sizes to assess the performance in both small and large trees. Moreover, we say a tree is *valid* if it does not become extinct and/or meets the criteria for the proportion of extant tips in either state, as described below. For trees simulated using slow rates, we use uniform sampling of rate parameters on the interval (0, 0.01]. For fast rates, we use the same distribution on the interval [0.1, 1].

**Table 1:** Summary of the simulation settings used to evaluate the robustness of the evolutionary scenario inference framework.

| Experiment | Description |
| --- | --- |
| complete-noreject | no missing taxa and no rejection for imbalanced tip states |
| missing-noreject | missing taxa and no rejection for imbalanced tip states |
| missing-noreject-sampling | missing taxa, no rejection for imbalanced tip states, and account for sampling proportions |
| complete-reject | no missing taxa and rejection for imbalanced tip states |
| missing-reject | missing taxa and rejection for imbalanced tip states |
| missing-reject-sampling | missing taxa, rejection for imbalanced tip states, and account for sampling proportions |

For simulations with missing taxa, we maintain the same observed extant-only tree sizes by simulating larger extant-only trees such that 20%, 40%, or 80% of the taxa are subsequently treated as missing. For each combination of observed tree size and missing-taxon proportion, we simulate 100 replicate trees. For simulations with tip state rejection, we reject trees in which either state contains fewer than 30% of the tips. For the experiments with sampling proportions, we assign the true proportions of missing states from simulations in the inference. Across all experiments, we initialize trees with root state *A*. Fixing the root state allows us to compare scenarios in which the root state corresponds to either the lower 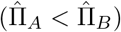 or higher 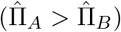 stationary frequency. We used the `tree.bisse` function from `diversitree` package (FitzJohn 2012) conditioned on tree size for the simulations.

For the inference procedure, we used `find.mle` function from the same package to perform MLE estimation on the rate parameters using `subplex` optimization method with a relative error tolerance of 10^−5^ and maximum number of iterations of 10000 per MLE search. We repeat the MLE search 10 times, each using an independent draw from the same probability distribution per parameter, for each tree.

To create a BiSSE likelihood function for each tree, we used the `make.bisse` function from the same package. For analyses using datasets with missing taxa, we set the argument `strict = F` to ignore cases where there is no variation in tip states. Using this option may affect the accuracy of the MLE estimation under BiSSE; however, this is unrelated to the purpose of this work, which is not to assess the performance of the BiSSE model, but rather to evaluate the statistical power of each evolutionary scenario being inferred. Moreover, for experiments that account for the sampling proportion of missing taxa in each state, we assigned a very small sampling proportion (*ρ* = 10^−7^) if the actual proportion missing in one state was 0, since the function does not allow a zero-valued proportion being assigned to either state.

Then, for each tree, we map both its true and estimated rates to a particular evolutionary scenario using Lemma 2, and compute the following probabilities for each experiment described in Table 1.

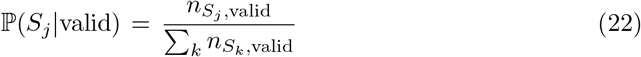

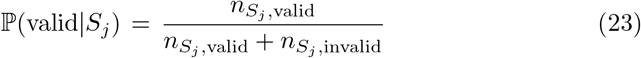

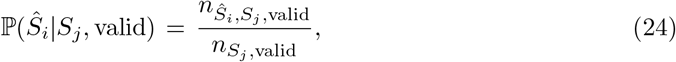

where *S*_*j*_ denotes true scenario *j* and *Ŝ*_*i*_ denotes estimated scenario 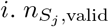 denotes the number of valid trees generated when the true scenario is 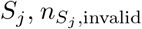 denotes the number of invalid trees generated when the true scenario *S*_*j*_, and 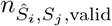 denotes the number of valid trees generated when the true scenario is *S*_*j*_ and the inferred scenario is *Ŝ*_*i*_.

Eq. (22) describes the relative prevalence of each true scenario among all valid trees, whereas Eq. (23) describes the probability that a tree generated under true scenario *S*_*j*_ is valid. Lastly, Eq. (24) describes the conditional probability of inferring a scenario *i* given that the true scenario is *j* and that the tree is valid. Using Eqs. (22) and (24) and knowing that each scenario from Lemma 2 is mutually exclusive, then using the Law of Total Probability, we can compute the following:

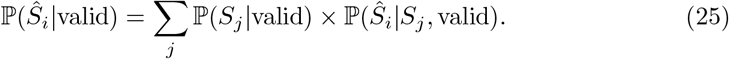

Eq. (25) describes the probability that a valid tree is inferred as evolutionary scenario *Ŝ*_*i*_. Next, for each true evolutionary scenario, we construct a confusion matrix for type-*I* (i.e., false positive) and type-*II* errors (i.e., false negative) between the true and estimated scenarios.

### 2.7 Application to real data

Here, we demonstrate how this concept of multiple evolutionary scenarios can be applied to real empirical data. We use the reconstructed timetree phylogeny of 6885 squamate species from Title et al. (2024), along with their trait data from SquamBase (Meiri 2024) for our analysis. Then, we sort the tree according to different Squamate families and infraorders, and only pick clades that have between 25 to 800 sampled extant species and run a BiSSE analysis on each clade. In total, we have clades of 38 families and 5 infraorders. Note that because taxonomic ranks are hierarchical, some family-level clades are nested within the infraorder-level clades included in the analysis. We therefore treat each clade as a separate empirical case rather than as statistically independent replicates. We also approximate the sampling proportion for each clade by comparing sampled species for each clade in the phylogeny with the list of species available on SquamBase. We also associated each species open versus closed habitat occupancy by recoded the Squambase dataset, with open habitats represented by: 1) Deserts and Xeric Shrublands, 2) Flooded Grasslands and Savannas, 3) Mangroves, 4) Mediterranean, 5) Montane Grasslands and Savannas, 6) Rock and Ice, 7) Temperate Grasslands, Savannas and Shrublands, and 8) Tropical and Subtropical Grasslands, Savannas and Shrublands, and closed habitats represented by: 1) Boreal Forests or Taiga, 2) Temperate Coniferous Forests, 3) Tropical and Subtropical Coniferous Forests, 4) Temperate Broadlead and Mixed Forests, 5) Tropical and Subtropical Dry Broadleaf Forests, and 6) Tropical and Subtropical Moist Broadleaf Forests. SquamBase associates each species with exactly one major biome type, meaning they can each be unambiguously assigned to either the open or closed state. For the inference procedure, we use `diversitree` package (FitzJohn 2012) and perform MLE estimation on the BiSSE rate parameters using `subplex` optimization method with a relative error tolerance of 10^−5^ and maximum number of iterations of 10000 per MLE search. We repeat the MLE search 50 times, each using an independent draw from the same probability distribution, namely on Uniform[0.0.1], per parameter, for each tree. We also obtained list of previously published BiSSE parameter estimates for a variety of angiosperm clades and traits, compiled by Helmstetter et al. (2023), to reinterpret using our new theoretical framework. As in the squamate dataset, some of the angiosperm clades are nested within others. Since we are only interested in studies that fitted fully unconstrained BiSSE models, we excluded cases where (1) the transition rate from one state to another was fixed at 0, (2) the transition rates were constrained to be equal in both directions, (3) the net diversification rates were constrained to be equal between the two states, or (4) the net diversification rate was fixed at 0 in either state. We also excluded studies that did not report their parameter estimates. In total, we report results based on 49 studies that met our selection criteria. Then, using the estimated rate parameters from each tree, we compute their expected stationary frequencies using the following formula (Maddison et al. 2007):

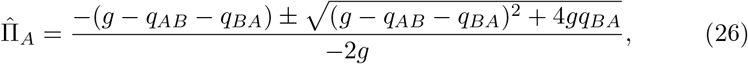

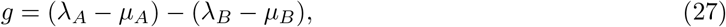

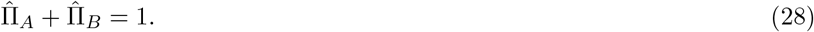

Then, for each study, we assign to a particular evolutionary scenario according to Lemma 2 and compute the proportion over all possible evolutionary scenarios.

## 3 Simulation and Inference Results

We report some key observations from the simulation experiments described in the methods section.

### 3.1 The number of rulesets is invariant among symmetrical stationary state frequencies

Using simulations, we show that the number of non-trivial rulesets is invariant among symmetrical stationary state frequencies, as claimed in Lemma 1 (Figs. S1-S2; Supplement). For BiSSE, all stationary frequency patterns in which both states have positive frequencies yield two non-trivial rulesets, whereas patterns in which either 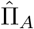 or 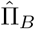 is zero yield no non-trivial rulesets. This is also established in Corollary 3. For GeoSSE, 8%, 12%, 50%, and 30% of the sampled stationary frequencies yield zero, one, two, and three non-trivial rulesets, respectively. Under GeoSSE, no non-trivial ruleset exists when (1) 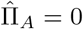, (2) 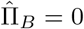, (3) 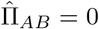, or (4) any combination of these conditions, assuming positive rates. No non-trivial ruleset also exists when 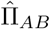 is equal to exactly one of 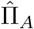 or 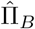, corresponding to the two diagonals through 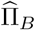 and 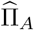, respectively, in Fig. S2. These findings also support Corollary S4.

### 3.2 Asymmetric realization of evolutionary scenarios as data-generating process

We compute ℙ (*S*_*j*_ | valid), as given in Eq. (22), among the set of valid trees under each simulation experiment described in Table 1. As seen in Table 2, scenarios 3a/3b are the most frequent, followed by 2a/2b, 4a/4b, and 1a/1b, respectively. Although the corresponding “a” and “b” scenarios are symmetric under exchange of the state labels by Definition 3 and Lemma 1, their observed proportions can differ because all simulations are initialized with the root in state *A*. This initialization breaks the state-label symmetry of the simulation design: for directionality class “a”, where 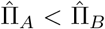, the process begins in the state with the lower stationary frequency, whereas for directionality class “b”, where 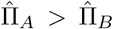, it begins in the state with the higher stationary frequency. Thus, differences between corresponding “a” and “b” directionality classes reflect, at least in part, the effect of the fixed initial (root) state rather than differences in their underlying rulesets. To validate this effect, we repeated the complete-noreject experiment with the root state randomly assigned to *A* or *B* with equal probability. Under this randomized initialization, the proportions of corresponding “a” and “b” directionality classes were more similar, consistent with the state-label symmetry of the underlying rulesets (Table S1).

**Table 2:**
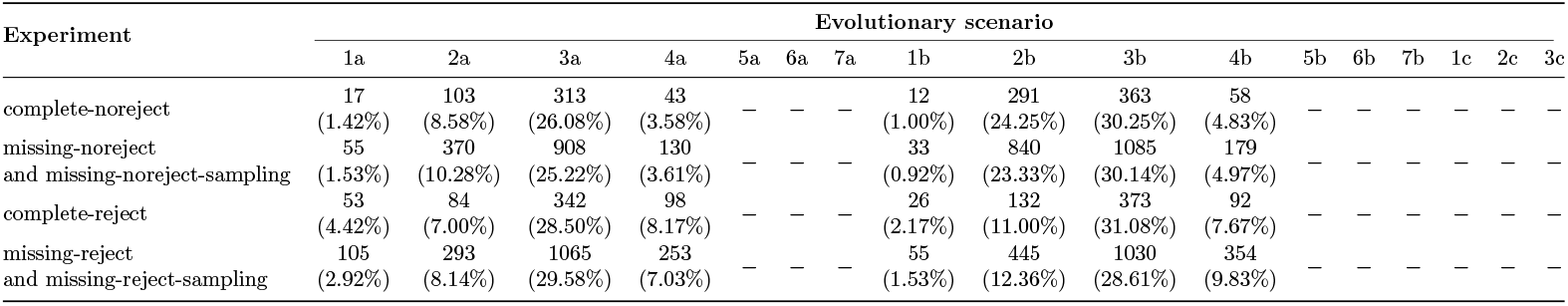
Distribution of true evolutionary scenarios among valid trees. For each simulation setting described in Table 1, the table reports the number and relative prevalence of each true scenario among all valid trees.

Both scenarios 3a and 3b require a net gain of species accumulation in both states (Lemma 2); thus, trees simulated under these scenarios are most likely to survive. In contrast, scenarios 5a/5b, 6a/6b, and 7a/7b correspond to cases involving either species loss in both states (scenarios 5 and 6) or equal rates of species accumulation in both states (scenario 7). The former is likely result in total extinction, whereas having exactly equal net diversification in both states is highly unlikely in the latter case. Similarly, scenarios 1c −3c correspond to the class of exactly equal state stationary frequencies in both states, which almost never occurs in real datasets. This is reflected in Table 2, where none of these scenarios are represented. Also, note that the proportion for each evolutionary scenario is fairly consistent across different simulation settings. Thus, the patterns observed above are not due to sampling bias.

Regarding the ability of each evolutionary scenario to generate valid trees, scenarios 4a/b are the most likely to produce valid trees, followed by scenarios 3a/b, 2a/b, and 1a/b, respectively, across all simulation settings (Table 1). By definition, scenarios 3a/b and 4a/b are source-source scenarios, in which both states have positive net diversification rates. Consequently, they are more likely to generate valid trees. In contrast, scenarios 1a/b and 2a/b are source-sink scenarios, where one state acts as a sink. Because the simulations are stochastic, lineages that enter the sink state are more likely to go extinct before speciating, reducing the probability of producing a valid tree. Similarly, scenarios 5a/b and 6a/b, in which both states are sinks, are less likely to generate valid trees. Lastly, scenarios 1c− 3c, corresponding to equal longterm tip state frequencies, and scenarios 7a/b with equal net diversification rates are also unlikely to be observed (Fig. 1).

**Fig. 1:**
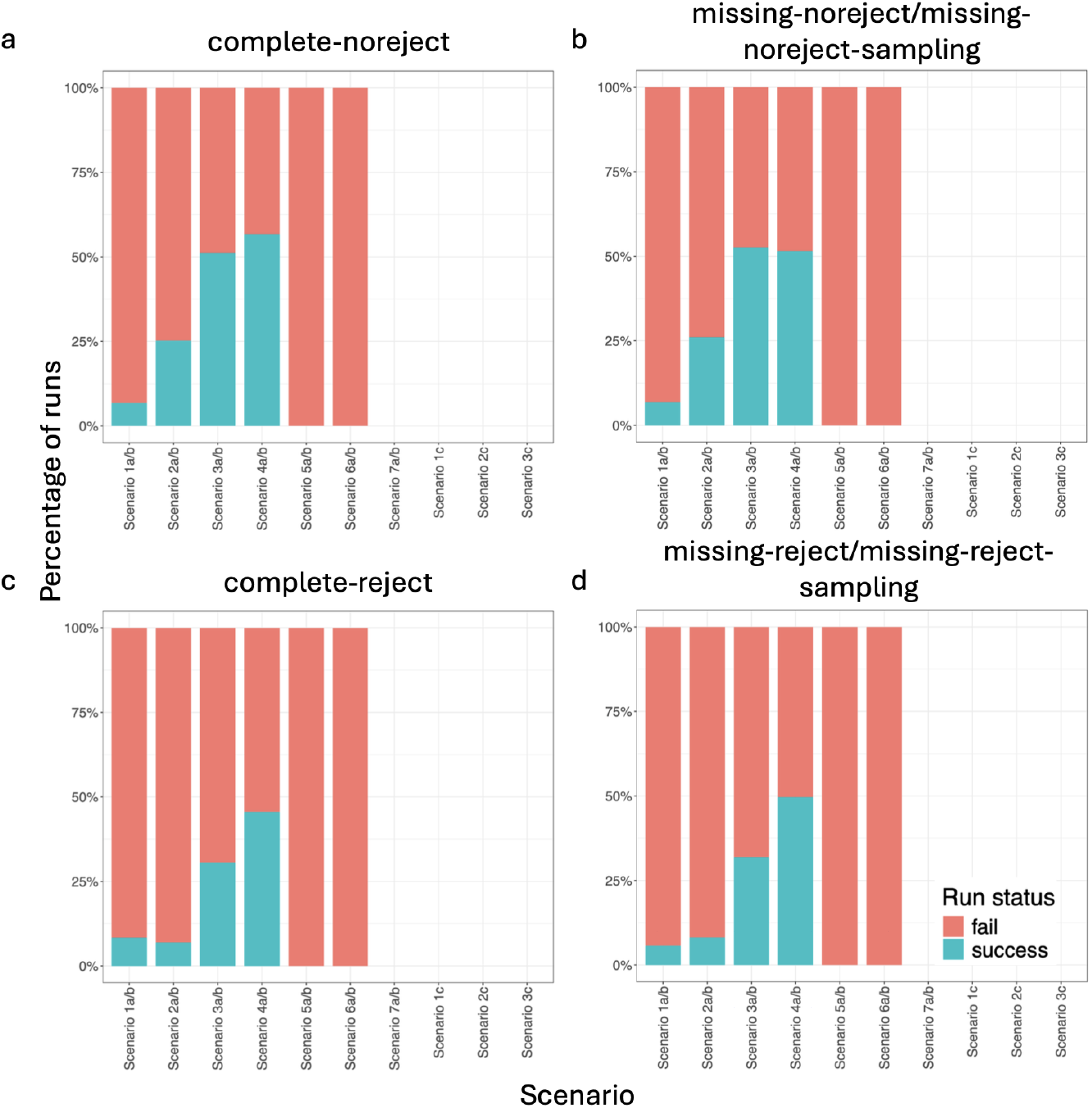
Probability of generating valid trees across evolutionary scenarios. Barplots show the proportion of successful (valid tree generated; blue) and failed (no valid tree generated; red) simulation runs for each evolutionary scenario under each simulation setting: (a) complete-noreject, (b) missing-noreject & missing-noreject-sampling, (c) complete-reject, and (d) missing-reject & missing-reject-sampling, as described in Table 1. The exact parameters and/or outcomes representing Scenarios 7a/b, 1c, 2c, and 3c were never simulated (blank bars).

### 3.3 Convergence toward stationary state frequencies between evolutionary scenarios

From Corollary 4, we expect that the distance from stationary state frequencies should decrease as the product of the spectral gap and the elapsed time of the process, *γt*, increases. Consistent with the theory, within each evolutionary scenario, the distance between the observed tip state frequencies and the stationary frequencies of the true generating process generally decreased with increasing *γt* (Fig. 2). This pattern is consistent among simulations using fast parameter rates and slow parameter rates cases (Figs. 2a-b).

**Fig. 2:**
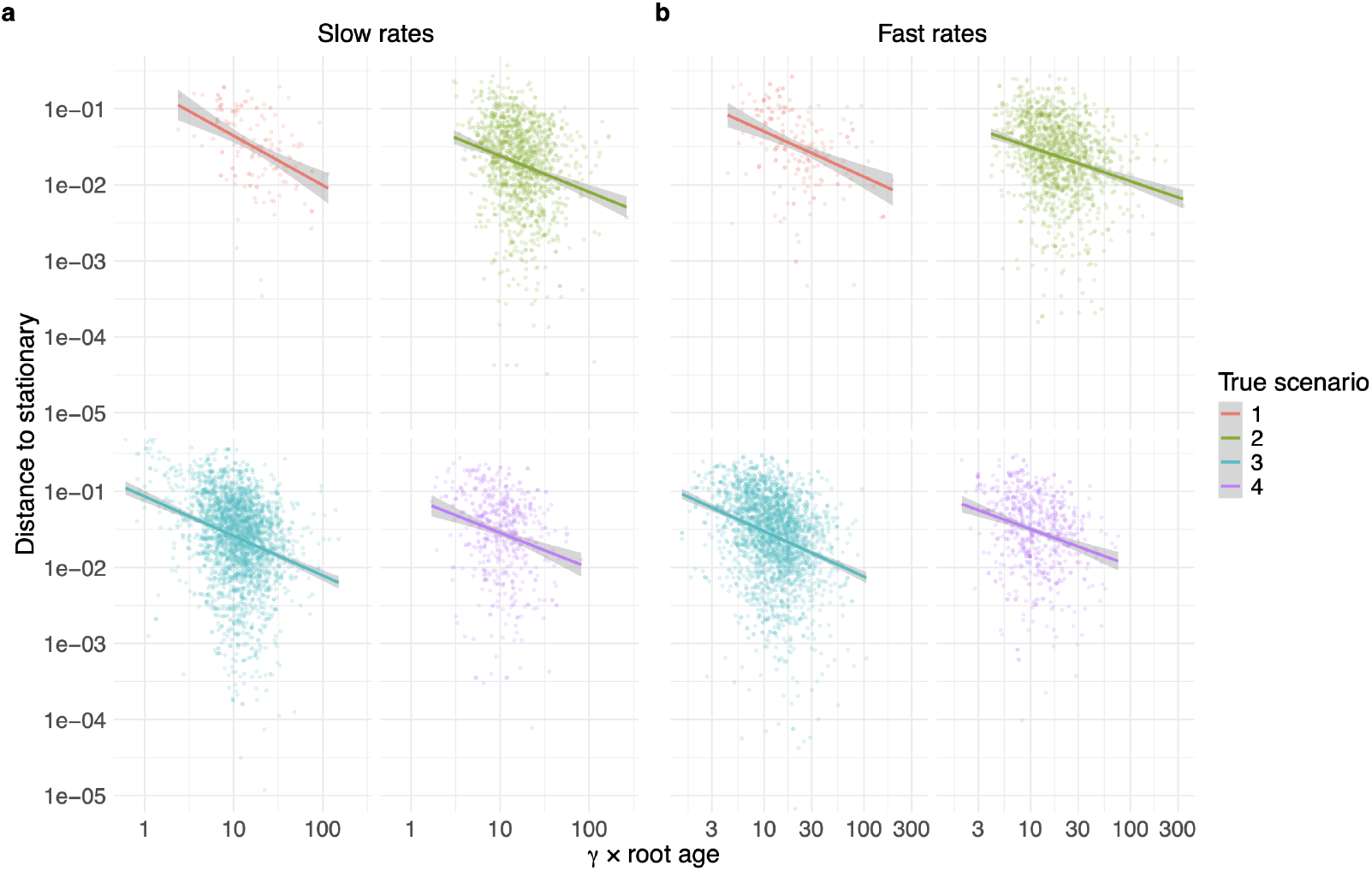
Relationship between the degree of convergence, measured by the product of the spectral gap (*γ*) and root age (*t*), *γt*, and the absolute distance between observed tip state frequencies and the stationary frequencies under the true generating process. Results are shown separately for datasets simulated using (a) slow and (b) fast parameter rates, and for true evolutionary scenarios 1–4 (combining the corresponding *a* and *b* scenarios). Points represent individual simulated trees, and lines show log–log linear regressions with 95% confidence intervals. Across all scenarios and both rate settings, larger values of *γt* are associated with smaller distances to stationarity, consistent with the expected relationship between the spectral gap and convergence toward stationary state frequencies.

### 3.4 Preferential inference of evolutionary scenarios using simulated datasets

Given datasets of extant-only trees, some evolutionary scenarios are inferred substantially more frequently than others. Across all simulation settings (Table 1), parameter estimation most frequently infers scenarios 3a/b, followed by scenarios 2a/b (Table 3), whereas scenarios 1a/b and 4a/b are less likely to be inferred. Interestingly, scenarios 3a/b also constitute the largest proportion of the true evolutionary scenarios among valid simulated trees in every experiment (Table 2).

**Table 3:** Probabilities of inferred evolutionary scenarios among valid trees under the different simulation settings described in Table 1. Each entry gives ℙ(*Ŝ*_*i*_| valid), corresponding to the probability that a randomly selected valid tree is inferred as evolutionary scenario *Ŝ*_*i*_.

| Estimated scenario | complete-noreject | missing-noreject | missing-noreject-sampling | complete-reject | missing-reject | missing-reject-sampling |
| --- | --- | --- | --- | --- | --- | --- |
| $\mathbb{P}(\hat{S}_{1a} \mid \text{valid})$ | 0.03 | 0.02 | 0.03 | 0.07 | 0.04 | 0.06 |
| $\mathbb{P}(\hat{S}_{2a} \mid \text{valid})$ | 0.15 | 0.13 | 0.17 | 0.13 | 0.11 | 0.15 |
| $\mathbb{P}(\hat{S}_{3a} \mid \text{valid})$ | 0.19 | 0.23 | 0.19 | 0.21 | 0.27 | 0.22 |
| $\mathbb{P}(\hat{S}_{4a} \mid \text{valid})$ | 0.02 | 0.02 | 0.02 | 0.05 | 0.05 | 0.05 |
| $\mathbb{P}(\hat{S}_{5a} \mid \text{valid})$ | 0 | 0 | 0 | 0 | 0 | 0 |
| $\mathbb{P}(\hat{S}_{6a} \mid \text{valid})$ | 0 | 0 | 0 | 0 | 0 | 0 |
| $\mathbb{P}(\hat{S}_{7a} \mid \text{valid})$ | 0 | 0 | 0 | 0 | 0 | 0 |
| $\mathbb{P}(\hat{S}_{1b} \mid \text{valid})$ | 0.04 | 0.03 | 0.04 | 0.07 | 0.05 | 0.06 |
| $\mathbb{P}(\hat{S}_{2b} \mid \text{valid})$ | 0.27 | 0.21 | 0.27 | 0.17 | 0.13 | 0.17 |
| $\mathbb{P}(\hat{S}_{3b} \mid \text{valid})$ | 0.26 | 0.31 | 0.24 | 0.22 | 0.29 | 0.23 |
| $\mathbb{P}(\hat{S}_{4b} \mid \text{valid})$ | 0.04 | 0.04 | 0.04 | 0.07 | 0.06 | 0.06 |
| $\mathbb{P}(\hat{S}_{5b} \mid \text{valid})$ | 0 | 0 | 0 | 0 | 0 | 0 |
| $\mathbb{P}(\hat{S}_{6b} \mid \text{valid})$ | 0 | 0 | 0 | 0 | 0 | 0 |
| $\mathbb{P}(\hat{S}_{7b} \mid \text{valid})$ | 0 | 0 | 0 | 0 | 0 | 0 |
| $\mathbb{P}(\hat{S}_{1c} \mid \text{valid})$ | 0 | 0 | 0 | 0 | 0 | 0 |
| $\mathbb{P}(\hat{S}_{2c} \mid \text{valid})$ | 0 | 0 | 0 | 0 | 0 | 0 |
| $\mathbb{P}(\hat{S}_{3c} \mid \text{valid})$ | 0 | 0 | 0 | 0 | 0 | 0 |

To investigate whether convergence toward stationarity may contribute to these differences, we compared the convergence properties of trees generated under each evolutionary scenario. Under simulations using fast and slow rates, trees generated under scenarios 1a/b tend to be the oldest, whereas scenarios 2a/b exhibit the largest spectral gaps (Figs. 3a-b; Figs. S3a-b). As a result, the scenarios differ in the product of the spectral gap and elapsed time (i.e., root age), *γt* (Fig. 3c; Fig. S3c). Consistent with Corollary 4, within each generating scenario, larger values of *γt* are associated with smaller distances between the observed tip state frequencies and the stationary frequencies under the true generating process (Fig. 2). Although the absolute distance to stationarity is broadly similar across evolutionary scenarios, scenarios 2a/b exhibit the smallest median distance despite scenarios 1a/b having the largest values of *γt* (Fig. 3d; Fig. S3d). Thus, although larger *γt* is associated with smaller distances to stationarity within each generating evolutionary scenario, differences in *γt* alone do not determine the relative distance to stationarity across scenarios, because the magnitude of the deviation also depends on properties of the underlying generating process.

**Fig. 3:**
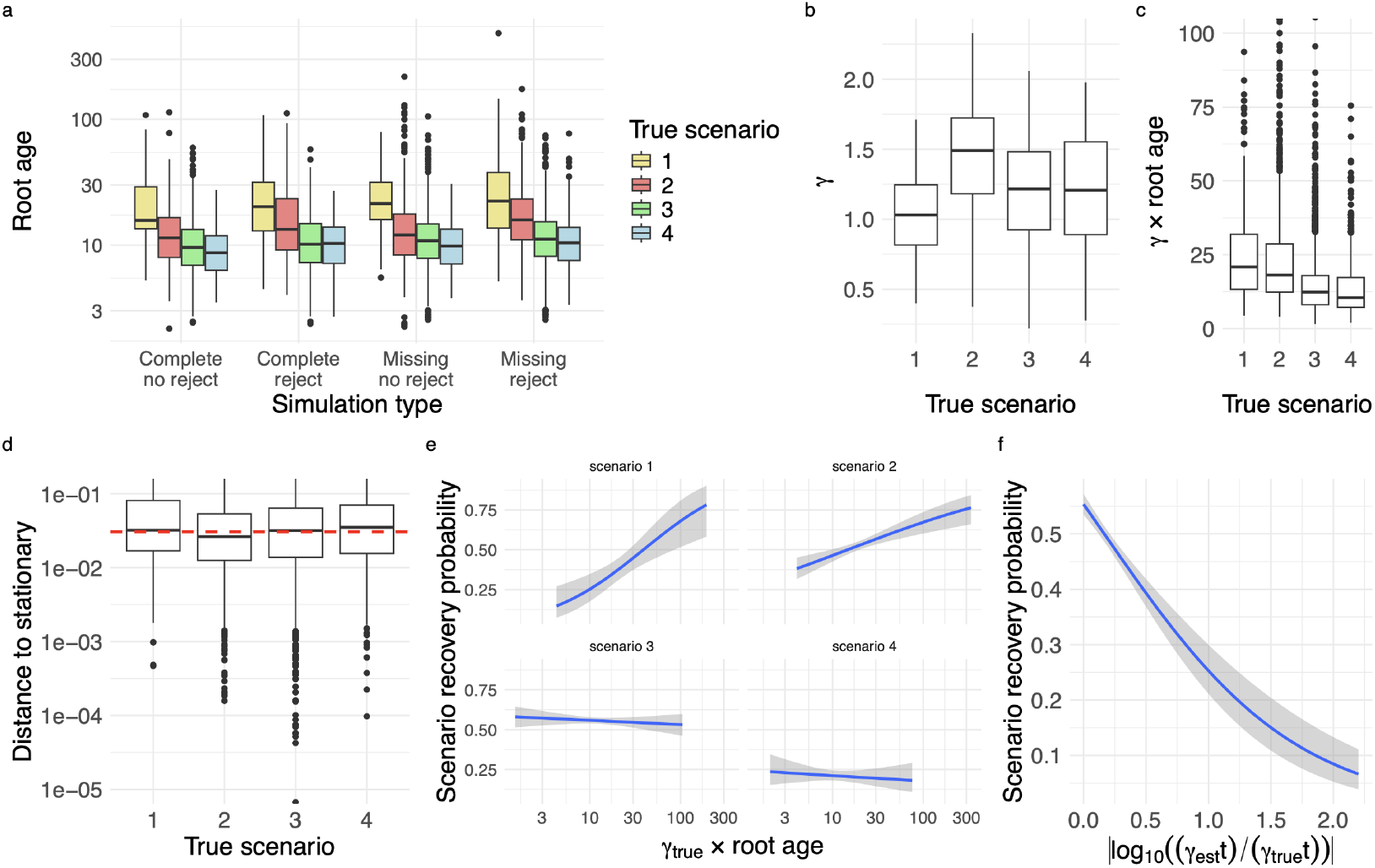
Convergence properties and evolutionary-scenario recovery for datasets simulated using fast parameter rates. (a) Distribution of root ages across the four simulation settings, as described in Table 1, grouped by true evolutionary scenario. (b) Distribution of the spectral gap, *γ*, computed using the true generating parameters for each evolutionary scenario. (c) Distribution of the product *γt*, where *t* denotes the root age. (d) Distance between the observed tip state frequencies and the stationary frequencies under the true generating process for each evolutionary scenario. (e) Probability of correctly recovering the true evolutionary scenario as a function of *γ*_true_*t*. Corresponding “a” and “b” directionality classes are pooled by scenario number for fitting and visualization, but recovery requires an exact match between the estimated and true evolutionary scenario, including the directionality class (e.g., 2a is correctly recovered only when inferred as 2a, not 2b). (f) Probability of correctly recovering the true evolutionary scenario as a function of the absolute log-ratio between the estimated and true spectral gaps, |log_10_(*γ*_est_*/γ*_true_) |. Lines in panels (e) and (f) show fitted binomial logistic regressions, with shaded regions indicating 95% confidence intervals. For all boxplots, center black lines indicate medians, boxes span the interquartile range (IQR), whiskers extend to the most extreme observations within 1.5*×* the IQR, and points beyond the whiskers indicate outliers.

We next examined whether convergence toward stationarity is associated with the probability of recovering the true evolutionary scenario. The relationship between *γt* and this probability differs substantially among generating evolutionary scenarios (Fig. 3e; Fig. S3e). Nevertheless, Scenario 2 consistently exhibited a positive relationship between *γt* and recovery probability under both fast and slow rates. For scenario 1, this relationship was strongly positive under fast rates but weaker under slow rates, whereas scenario 3 showed a weak negative relationship under fast rates but a positive relationship under slow rates. In contrast, scenario 4 exhibited a negative relationship under both rate regimes. Thus, greater convergence toward stationarity does not universally improve scenario recovery. However, the consistent positive relationship for scenario 2 suggests that convergence may partly explain its relatively high recovery probability and, consequently, may explain the relatively frequent inference of scenarios 2a/b in Table 3. In contrast, the frequent inference of scenarios 3a/b cannot be explained by convergence alone, because the relationship between *γt* and recovery probability is inconsistent across the two rate regimes. In addition, the probability of recovering the true scenario decreases substantially as the difference between the estimated and true spectral gaps increases (Fig. 3f; Fig. S3f). This indicates that accurate estimation of the convergence rate is strongly associated with correct scenario recovery.

The remaining scenarios—5a/b, 6a/b, 7a/b, and 1–3c—are not inferred at all (Table 3). This result is reassuring, as these scenarios are also unlikely to represent the true evolutionary processes, given that they rarely generate valid trees (Table 2).

### 3.5 Uneven sensitivity across evolutionary scenarios

Across all the experiments described in Table 1, the scenarios 1a/b and 4a/b tend to have a high rate for producing type −*II* error (false negative) (Table 4; Tables S4–S8). In other words, BiSSE frequently fails to recover these scenarios when they are the true generating scenarios, indicating low sensitivity. Consistent with this result, Table 3 also shows that scenarios 1a/b and 4a/b are inferred less frequently than the other scenarios. Conversely, scenarios 1a/b and 4a/b generally have relatively low type −*I* error rates (false positive), whereas scenarios 2a/b and 3a/b have higher false positive rates (Table 4; Tables S4–S8). Thus, trees generated under other scenarios are more frequently misclassified as scenarios 2a/b and 3a/b than as scenarios 1a/b and 4a/b. This asymmetry in both sensitivity and false positive rates may therefore contribute to the greater overall frequency with which scenarios 2a/b and 3a/b are inferred in Table 3.

**Table 4:** Prediction accuracy for each true scenario versus all other scenarios using the dataset simulated under the complete-noreject setting described in Table 1. Scenarios 5a/b, 6a/b, 7a/b, 1c, 2c, and 3c are omitted because they have no valid-tree representation.

|  |  | Predicted scenario |  |
| --- | --- | --- | --- |
|  |  | 1a | Other scenarios |
| True scenario | 1a | 0.29 | 0.71 |
|  | Other scenarios | 0.03 | 0.97 |
|  |  | 2a | Other scenarios |
|  | 2a | 0.61 | 0.39 |
|  | Other scenarios | 0.10 | 0.90 |
|  |  | 3a | Other scenarios |
|  | 3a | 0.52 | 0.48 |
|  | Other scenarios | 0.07 | 0.93 |
|  |  | 4a | Other scenarios |
|  | 4a | 0.21 | 0.79 |
|  | Other scenarios | 0.02 | 0.98 |
|  |  | 1b | Other scenarios |
|  | 1b | 0.25 | 0.75 |
|  | Other scenarios | 0.04 | 0.96 |
|  |  | 2b | Other scenarios |
|  | 2b | 0.58 | 0.42 |
|  | Other scenarios | 0.17 | 0.83 |
|  |  | 3b | Other scenarios |
|  | 3b | 0.50 | 0.50 |
|  | Other scenarios | 0.16 | 0.84 |
|  |  | 4b | Other scenarios |
|  | 4b | 0.22 | 0.78 |
|  | Other scenarios | 0.03 | 0.97 |

### 3.6 Preferential inference of evolutionary scenarios using empirical datasets

As shown in Fig. 4, BiSSE most frequently infers scenarios 3a/b for squamate clades, consistent with the simulation results (Table 3). Compared with the simulations, however, the empirical analysis infers scenarios 4a/b more frequently and scenarios 2a/b less frequently (Fig. 4a). Past studies that used BiSSE on Angiosperm datasets also indicate that scenarios 3a/b and 4a/b are most frequently inferred (Fig. 4b). Note that we could not perform the corresponding analyses of the spectral gap, the product of the spectral gap and root age, or distance to stationarity for the angiosperm datasets because the information required for these calculations, including root ages and observed tip state frequencies, was not consistently reported across studies.

**Fig. 4:**
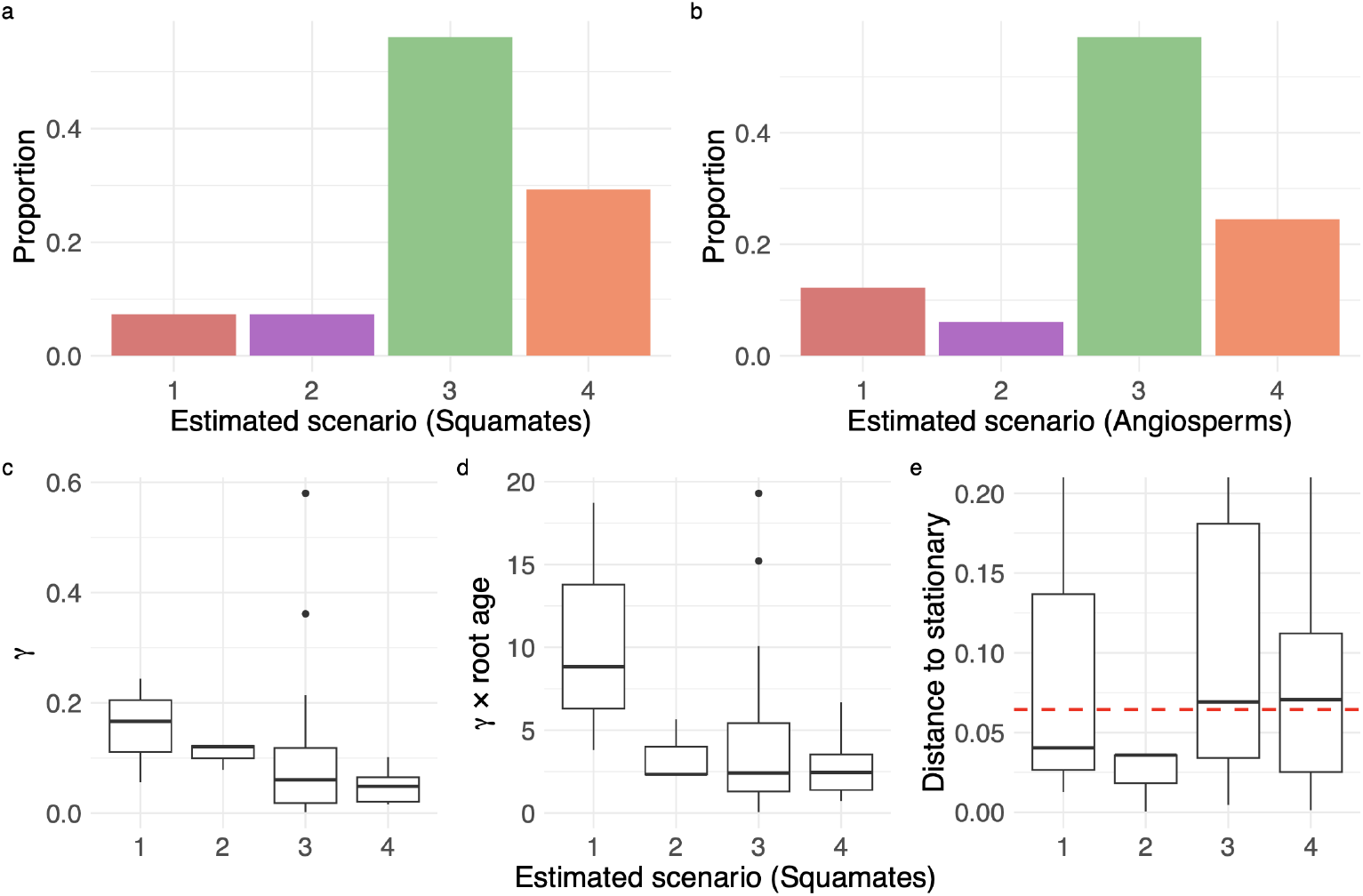
Empirical analysis under the BiSSE model. Proportion of empirical clades assigned to each evolutionary scenario for (a) squamates and (b) angiosperms. For the squamate dataset, distributions across estimated scenarios of (c) the spectral gap, *γ*, computed using the estimated parameters, (d) the product of the estimated spectral gap and root age, *γt*, and (e) the distance between observed tip state frequencies and their corresponding expected stationary frequencies. The dashed red line in (e) indicates the median distance to stationarity across all empirical squamate clades.

One possible explanation for this difference is that clades inferred as scenarios 3a/b or 4a/b have distributions of absolute distances between the observed tip state frequencies and their expected stationary frequencies that more closely resemble the overall distribution across all empirical trees than do clades inferred as scenarios 1a/b or 2a/b (Fig. 4e). In contrast, both scenarios 1a/b and 2a/b have similar distributions of absolute distances to their expected stationarity, with both scenarios tend to be closer to their stationary expectations than the empirical squamate dataset as a whole. If we assume that the estimated scenarios reasonably approximate the true underlying evolutionary processes, this pattern suggests that empirical clades assigned to scenarios 3a/b or 4a/b span distances to stationarity that are more representative of the empirical dataset as a whole, whereas clades assigned to scenarios 1a/b or 2a/b tend to be substantially closer to their stationary expectations. This pattern is less pronounced in the simulation results, where distances to stationarity calculated from the true parameter values overlap substantially across the four true scenarios, and the median distance for scenarios 2a/b remains relatively close to the overall median across simulated trees (Fig. 3d; Fig. S3d).

## 4 Discussion

In this study, we have established formal functional relationships (“rulesets”) between rate parameters and the expected long-term tip state frequencies produced by branching processes under the state-dependent speciation and extinction (SSE) model class. As mathematical objects, each ruleset defines a distinct, non-overlapping solution space consisting of a set of equalities and inequalities among model parameters that is compatible with a particular pattern of long-term stationary frequencies (defined as proportions) across character states. As biological objects, each ruleset corresponds to a particular evolutionary scenario, describing the relationships among the rates at which species originate, go extinct, and transition among discrete states.

This study examines the relationship between tip state frequencies and the evolutionary processes that generate them by asking a simple question: in the absence of a phylogenetic tree, how much information about the underlying evolutionary process can be extracted from the tip state frequencies of extant species alone? This question is relevant because biologists are often interested in determining which evolutionary scenarios provide plausible explanations for observed patterns of character states. Such scenarios are typically inferred using a phylogenetic tree that represents the evolutionary relationships among the species in the dataset. However, phylogenetic reconstruction depends on the availability and quality of sequence data, as well as the assumption that evolutionary history can be adequately represented by a tree. Our framework is not intended to replace tree-based inference. Rather, it provides a complementary approach that asks what can be learned from a minimal amount of information—namely, the observed species and their character states even when their phylogenetic relationships are not available. By identifying the alternative rulesets compatible with a given pattern of tip state frequencies, our framework characterizes the range of evolutionary scenarios that remain plausible given this minimal information.

Moreover, we can further classify these rulesets according to their triviality. We consider a ruleset trivial if the net diversification rates across all states belong to the same dynamical phase: either all are supercritical (source; *ρ >* 0, *λ*_*i*_ *> µ*_*i*_ for all *i*) or all are subcritical (sink; *ρ <* 0, *λ*_*i*_ *< µ*_*i*_ for all *i*). In contrast, non-trivial rulesets are those where their states in opposing dynamical phases, with some states supercritical and others subcritical. Within each directionality class of stationary tip state frequencies (e.g., 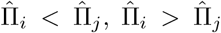, or 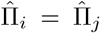), we establish a bijection between rulesets and evolutionary scenarios under a given SSE model. Importantly, the same long-term tip state frequencies can be compatible with multiple distinct rulesets and, consequently, multiple evolutionary scenarios. Furthermore, each ruleset defines a solution space containing infinitely many parameter combinations that yield the same stationary frequencies. As a proof of concept, we demonstrate these results using the BiSSE and GeoSSE frameworks, with additional GeoSSE results provided in the Supplement. However, the concepts developed here can be extended to other discrete-state SSE models.

There are two complementary perspectives for examining how the data-generating process governs lineage diversification: forward-in-time, by asking which evolutionary scenarios are more likely to generate valid trees, and backward-in-time, by asking which scenarios are more likely to be inferred from reconstructed trees and their observed tip states. Under the forward-in-time perspective, because each evolutionary scenario is associated with a unique ruleset and each ruleset occupies a distinct, non-overlapping region of the parameter space, we can quantify the probability that each ruleset generates valid trees. This allows us to assess which rulesets are more likely to be realized during the tree-building process for a given model. Through simulations, we found that although the same long-term frequency can be generated under multiple distinct evolutionary scenarios, these scenarios are not all equal in terms of generating valid trees. By valid trees, we refer to trees that leave sampled descendants at the present and/or meet a pre-defined degree of tip state imbalance. Scenarios where both states are source, such as scenarios 3a/b and 4a/b, are more likely to generate valid trees, followed by scenarios where only one of the states is source, such as scenarios 1a/b and 2a/b. Scenarios where both states are sink, such as scenarios 5a/b and 6a/b, are unlikely to produce valid trees (Fig. 1). Corollary 2 shows that scenarios 3a/b and 4a/b correspond to supercritical branching processes (*ρ >* 0), whose expected lineage abundances grow exponentially over time. Consequently, they are more likely to generate large, non-extinct trees. Conversely, scenarios 5a/b and 6a/b correspond to subcritical branching processes (*ρ <* 0), whose expected lineage abundances decline over time, increasing the probability of extinction.

When examining the relative prevalence of each true scenario among all valid trees, scenarios 4a/b are substantially less represented (approximately 3–4 times less frequent) than scenarios 3a/b, despite both corresponding to supercritical branching processes (*ρ >* 0) and positive net diversification rates in both states (Table 2). We think this difference arises because scenarios 4a/b require transition rates that oppose the ordering of their stationary tip state frequencies within each directionality class. For example, scenario 4a has a higher transition rate into state A than into state B, while remaining in the directionality class where 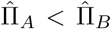. Consequently, the net diversification rate in state B must be sufficiently greater than that in state A to maintain the prescribed stationary frequencies (Table S2). In contrast, scenarios 3a/b require transition rates that are consistent with the ordering of their stationary frequencies. Therefore, the admissible parameter space for scenarios 4a/b is more restricted than that for scenarios 3a/b, making them less likely to be realized during the tree-generation process. Scenarios 2a/b are also more prevalent among valid trees than scenarios 1a/b, despite both corresponding to critical branching processes *ρ* = 0 (Table 2). This difference might arise because scenarios 1a/b require the transition rate into the sink state to compensate for the net loss of species in that state while remaining within the prescribed stationary class (Table S2). In contrast, as shown in Lemma 2, scenarios 2a/b impose no constraints on the relative magnitudes of the two transition rates to remain within their respective directionality classes. These patterns are consistent across all simulation settings, including complete sampling, missing taxa with and without accounting for sampling proportions, and simulations with both low and high variation in tip state frequencies. Scenarios 5a/b, 6a/b, 7a/b, and 1c − 3c failed to produce a single representative in these experiments (Table 2). Scenarios 5a/b and 6a/b correspond to subcritical branching processes, and are therefore prone to extinction. In contrast, scenarios 7a/b require equal net diversification rates in both states, which is highly unlikely under continuous random sampling of the model parameters. Similarly, scenarios 1c − 3c correspond to equal stationary tip state frequencies in both states, which is also unlikely to arise through random parameter sampling.

These forward-in-time results may help inform priors in tree-based Bayesian inference by identifying parameter regimes capable of generating trees consistent with the observed data. Because empirical datasets typically consist of extant-only trees conditioned on survival, supercritical branching processes (e.g., scenarios 3 and 4) are naturally more likely to generate such datasets. Furthermore, the choice between scenarios 3 and 4 may depend on prior biological knowledge regarding the role of anagenetic transitions. For example, one may prefer scenarios in which the inferred direction of state transitions is consistent with prior expectations about trait evolution. In contrast, for systems expected to remain near equilibrium, such as diversity-dependent diversification models (Soewongsono and Landis 2026), priors based on scenarios derived under critical processes (*ρ* = 0) may be more appropriate. Because evolutionary scenarios and rulesets are bijective within each directionality class, priors on the model parameters can also be constructed using the functional relationships among rate parameters implied by these rulesets. Lastly, although empirical phylogenies provide only present-day tip state frequencies, our framework remains applicable because these observations can be interpreted as snapshots of an underlying branching process that may be approaching its asymptotic tip state frequencies.

From the backward-in-time perspective, we next asked whether this asymmetry persists in inference: given only a present-day reconstructed phylogeny (i.e., with extinct lineages pruned) and the character states observed at the tips, are some evolutionary scenarios more likely to be inferred than others? Through simulation experiments, we show that this is indeed the case. In particular, scenarios 3a/b are the most favored, followed by scenarios 2a/b. In all cases, scenarios 1a/b and 4a/b share a similar probability of being chosen as the preferred scenario (Table 3). The frequent inference of scenarios 3a/b is consistent with their prevalence among valid simulated trees and may partly reflect conditioning of reconstructed-tree likelihoods on survival. Because scenarios 3a/b correspond to supercritical source-source dynamics (*ρ >* 0), they are more likely to produce non-extinct trees that survive to the present. This tendency is relevant because reconstructed-tree likelihoods are commonly conditioned on survival of the lineages descending from the root. Similar effects may arise when conditioning on the number of extant taxa or the elapsed time of the process, because supercritical processes are more likely to generate large trees and to persist over longer periods without extinction.

Interestingly, scenarios 4a/b, which also represent supercritical processes in both states, are inferred less frequently than 3a/b (~ 5 times smaller; Table 3). One possible explanation is that scenarios 4a/b require a competing balance between lineage diversification and anagenetic transitions to remain within their corresponding stationary frequency class. For example, under Scenario 4a, speciation rate in state B must exceed the transition rate out of state B. In addition, because trees are observed at a finite time (e.g., the present) rather than at equilibrium, the likelihood may favor scenarios that better reconcile the observed distance to stationarity with their expected rate of convergence. For example, a younger tree that is already near stationarity may be better explained by a scenario with a faster rate of convergence. Another contributing factor may be misclassification among scenarios. Across simulation settings, scenarios 2a/b and 3a/b generally have false positive rates approximately 2–4 times those of scenarios 1a/b and 4a/b (Table 4; Tables S4–S8). Thus, the frequent inference of scenarios 3a/b may partly reflect misclassification of trees generated under other scenarios as 3a/b.

In the simulated datasets, simpler source-sink dynamics such as scenarios 2a/b are more frequently inferred than scenarios 4a/b (Table 3). Parameters drawn under scenarios 2a/b generally correspond to faster rates of convergence to the stationary distribution than those drawn under scenarios 4a/b (Fig. 3b; Fig. S3b). Likewise, the product of the convergence rate and elapsed time, *γt*, tends to be larger under scenarios 2a/b than under scenarios 4a/b. Theoretically, for a given process, larger values of *γt* imply greater convergence toward the stationary distribution over the observed lifetime of the process.

Consistent with this expectation, trees generated under scenarios 2a/b tend to have smaller distances to stationarity than those generated under scenarios 4a/b, although their distributions overlap substantially (Fig. 3d; Fig. S3d). Moreover, under scenarios 2a/b, larger values of *γt* are consistently associated with a higher probability of recovering the true scenario (Fig. 3e; Fig. S3e). More generally, the probability of recovering the true evolutionary scenario decreases as the discrepancy between the estimated and true spectral gaps increases (Fig. 3f; Fig. S3f), suggesting that accurately capturing the timescale of convergence toward the stationary distribution is important for recovering the underlying evolutionary scenario. Together, these results suggest that faster convergence toward stationary frequencies may partly contribute to the greater recoverability of scenarios 2a/b relative to scenarios 4a/b, although convergence alone does not explain the differences in their overall inference frequencies. As with scenarios 3a/b, the frequent inference for scenarios 2a/b may also partly reflect misclassification, as trees generated under other scenarios are more frequently misclassified as scenarios 2a/b than as scenarios 1a/b or 4a/b (Table 4; Tables S4–S8). Thus, supercriticality alone does not explain why some evolutionary scenarios are inferred more frequently than others; the observed differences also appear to reflect convergence toward stationary frequencies, scenario-specific sensitivity and false positive rates, and the structure of the parameter space. In particular, scenarios 1a/b and 4a/b have relatively low sensitivity and are therefore less likely to be recovered when they are the true generating scenarios. Lastly, scenarios corresponding to identical stationary frequencies in both states (1c–3c) are expected to be extremely rare in both forward-in-time simulation and backward-in-time inference (Tables 2 and 3).

When applying BiSSE to empirical data, including clades of infraorders and families from the reconstructed squamate phylogeny and previously published BiSSE analyses of angiosperm clades and traits, we found that scenarios 3a/b are the most frequently inferred scenarios in both systems (Fig. 4a,b). This pattern is consistent with our simulation results. One possible contributing factor is that both the empirical phylogenies and the likelihood calculations are based on reconstructed trees containing lineages that survived to the present. Such conditioning may favor evolutionary processes that are more likely to produce surviving reconstructed trees, including supercritical dynamics such as scenarios 3a/b and 4a/b.

In contrast to the simulation results, the empirical analyses infer scenarios 4a/b more frequently than scenarios 2a/b (Fig. 4a,b). For the squamate dataset, parameter estimates associated with scenarios 2a/b correspond to larger spectral gaps than those associated with scenarios 4a/b (Fig. 4c), indicating faster convergence towards stationarity. However, clades assigned to scenarios 2a/b tend to be younger than those assigned to scenarios 4a/b. Consequently, the distributions of *γt* for the two scenarios are relatively similar (Fig. 4d). Despite this similarity, clades assigned to scenarios 4a/b tend to exhibit larger distances between their observed tip state frequencies and the expected stationary frequencies than those assigned to scenarios 2a/b (Fig. 4e). This is consistent with our simulation results showing that, although *γt* describes the rate of convergence over the lifetime of a given process, differences in *γt* alone do not determine absolute distances to stationarity across different evolutionary processes.

We caution that these empirical patterns should be interpreted carefully because they assume that the estimated scenario adequately represents the true underlying evolutionary scenario for each dataset. The relatively high frequency of scenarios 3a/b and 4a/b could also reflect model misspecification. BiSSE is known to be susceptible to elevated type −*I* error when hidden states affecting lineage diversification are not considered (Rabosky and Goldberg 2015; Beaulieu and O’Meara 2016). More generally, processes not represented by the fitted BiSSE model could influence both parameter estimates and the evolutionary scenarios to which empirical clades are assigned. In addition to model misspecification, the choice of prior distributions for model parameters may influence the relative support for different scenarios.

Overall, our results show that distinct data-generating processes can produce identical long-term tip state frequencies while corresponding to distinct evolutionary scenarios. Nevertheless, because phylogenetic trees are reconstructed using only extant species and likelihood calculations are performed on these reconstructed trees, inference tends to favor a limited subset of evolutionary scenarios. In particular, scenarios corresponding to supercritical processes (i.e., source-source dynamics) are more frequently inferred. This preference arising from conditioning on reconstructed trees should be distinguished from statistical biases previously identified in SSE models. For example, inference under SSE models can be more reliable when applied to sufficiently large trees with relatively balanced distributions of tip states or when hidden states affecting diversification are accounted for (Davis et al. 2013; Gamisch 2016).

Furthermore, by relating the spectral gap (convergence rate), determined by combinations of model parameters, to the expected stationary state frequencies generated by those parameters, our framework provides additional information about how evolutionary processes approach their long-term behavior. These relationships may help guide the construction of biologically and dynamically informed prior expectations and identify parameter combinations that are inconsistent with observed patterns of tip state frequencies. More broadly, both the concepts and theoretical framework developed in this study can be extended to other SSE models, although such extensions can become increasingly complex as the state space grows. For example, in a GeoSSE model with three regions *A, B*, and *C*, directionality classes could be defined either across the full set of geographic range states (*A, B, C, AB, AC, BC, ABC*) or at the region level (*A, B, C*). The number of possible directionality classes increases substantially with the number of states because of the many possible equality and inequality relationships among their stationary frequencies. Developing efficient approaches for characterizing these higher-dimensional directionality classes and their associated rulesets therefore represents an important direction for extending the framework to more complex SSE models, providing a foundation for understanding how increasingly complex data-generating processes shape the stationary patterns observed across character states.

## Acknowledgments

The authors thank all members of the Landis lab at Washington University in St. Louis for their constructive feedback on this manuscript. We also thank the anonymous reviewers for their careful reading and suggestions, which strengthened this work.

## Declarations

### Code availability

All relevant scripts used in this study are publicly available at https://github.com/alberts2/diffusion rate pattern SSE.

### Author contributions

A.C.S. and M.J.L designed research; performed research; analyzed data; and wrote the manuscript.

## Supplementary Materials

### 1.1 Proof of Theorem 1

Let *N*_*i*_(*t*) denote the number of extant lineages in state *i* at time *t*. Recall that 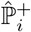 and 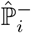 denote the probability of gaining or losing a new species in state *i*. At stationarity, the expected number of events that increase the number of lineages in state *i* must balance the expected number of events that decrease it. Thus:

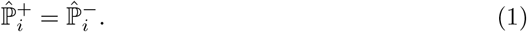

Next, because Π_*i*_(0) denotes the initial frequency of species in state *i*, the collection *{*Π_*i*_(0)*}*_∀*i*∈*S*_ must form a valid probability distribution. Hence,

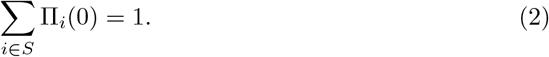

with Π_*i*_(0) ≥ 0, ∀ *i* ∈ *S*. Likewise, the components of the speciation, extinction, and anagenetic transition-rate vectors represent event rates and are therefore restricted to the admissible positive parameter space.

To distinguish evolutionary rulesets associated with a given set of stationary frequencies, we additionally impose pairwise consistency between the ordering of the stationary frequencies and the corresponding total net rates. Thus, for every pair of states, 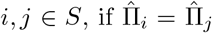, we require the corresponding states are required to have equal total net rates, i.e. Φ_*total,i*_ = Φ_*total,j*_. Similar requirements exist for the greater than and less than cases. Applying this condition to every pair *i, j* ∈ *S* ensures that the ordering of the total net rates is consistent with the ordering of the given stationary frequencies and therefore defines the class of distinct evolutionary rulesets considered here. Hence, any parameter set satisfying the stationary balance equation and parameter-space constraints, together with the imposed pairwise ordering criterion, constitutes an admissible evolutionary ruleset associated with the given stationary frequencies. □

### 1.2 Proof of Corollary 1

Recall that 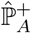 and 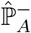 denote the probability of gaining or losing a new species in state *A* under stationary scenario, respectively. In the BiSSE model, we can gain a new species in *A* through a speciation event acting on a parent species in state *A* and through an anagenetic change from state *B* to state *A*. Therefore:

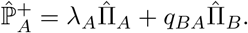

On the other hand, we lose a new species in *B* through an extinction event for a species in state *B* and through an anagenetic change from state *A* to state *B*. Therefore:

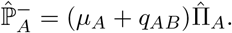

Similarly, we can define:

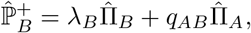

and

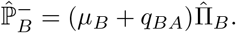

Assuming that there is no change in expected abundance of species in each state, we have:

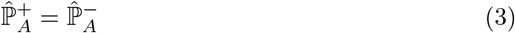

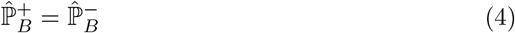

Φ_*total,A*_ and Φ_*total,B*_ represent the flux between all incoming and outgoing rates from events occurring in state *A* and *B*, respectively. In BiSSE, these come from the rates of speciation, extinction, and anagenetic change. Thus:

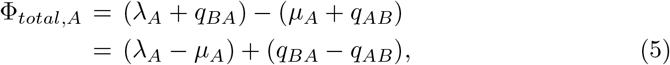

and

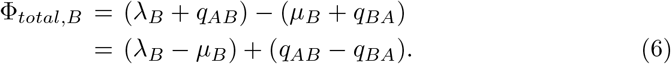

From Eqs (3) and (4), we have:

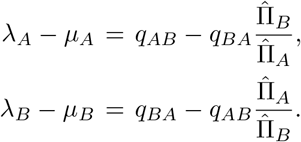

Substituted into Eqs. (5) and (6) we have:

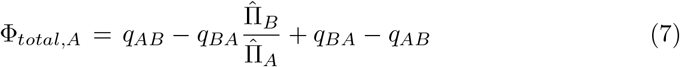

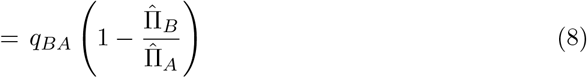

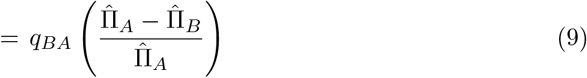

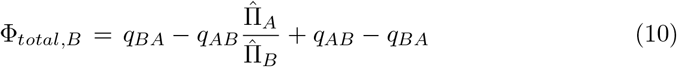

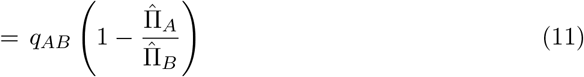

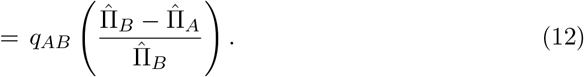

The relationship between Φ_*total,A*_ and Φ_*total,B*_ follows directly from the relationship between 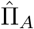 and 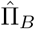.

Furthermore, from Eqs. (3) and (4) we have:

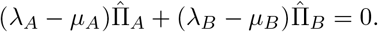

For BiSSE models where there is no expected change in abundance of species in each state, when both stationary state frequencies are positive, 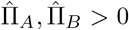, and both net diversification rates are nonzero, *λ*_*i*_ ≠ *µ*_*i*_, ∀ *i* ∈ *S*, the equation above requires the net diversification rates in states *A* and *B* to have opposite signs. Therefore, the net diversification rates of the two states cannot be equal. □

### 1.3 Proof of Corollary 2

The proof is similar to the proof of Corollary 1. However, here, instead of assuming zero net change in expected abundance in both states, we assume both states to grow at the equal rate *ρ*. Specifically,

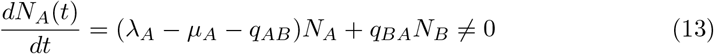

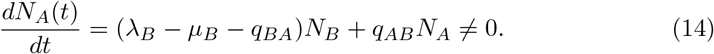

Since 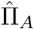 and 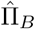 are stationary frequencies in *A* and *B*, then the following is true as *t* → ∞:

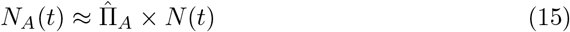

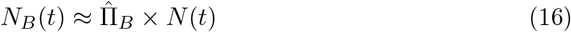

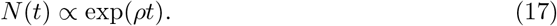

Thus, using Eq. (17) and substituting both Eqs. (15) and (16) into Eqs. (13) and (14) after differentiating them w.r.t. *t*, we obtain:

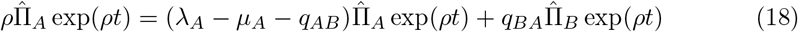

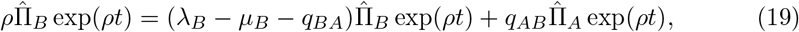

which implies:

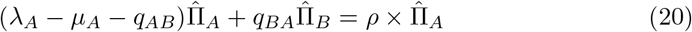

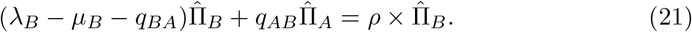

Next, choosing 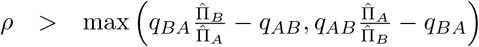 ensures that *λ > µ*_*A*_ and *λ*_*B*_ *> µ*_*B*_ (i.e., both states are source). Similarly, choosing 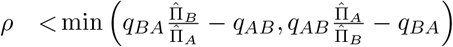 ensures that *λ*_*A*_ *< µ*_*A*_ and *λ*_*B*_ *< µ*_*B*_ (i.e., both states are sink). □

### 1.4 Rulesets in a GeoSSE model

We show how to derive a ruleset given a set of stationary frequencies of species count in different ranges in the context of a GeoSSE model (Goldberg et al. 2011). For simplicity, we describe in the case of *n* = 2 regions in Corollary S1.

#### Corollary S1.

*Given a GeoSSE with state space S* = *{A, B, AB*}, *set of stationary frequencies*, 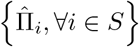, *and initial state frequencies* {Π_*A*_(0), Π_*B*_(0), Π_*AB*_(0) : Π_*A*_(0) + Π_*B*_(0) + Π_*AB*_(0) = 1}. *We obtain a ruleset(s) by solving the following system of equations:*

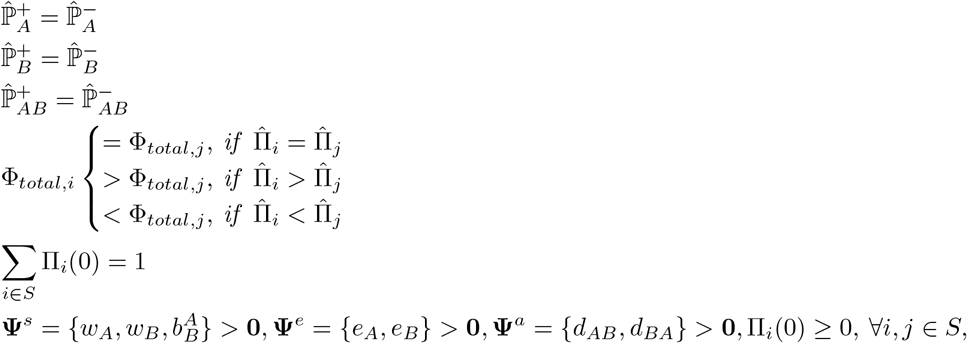

*where w*_*A*_ *and w*_*B*_ *are within-region speciation rates in region A and B, respectively*. 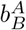 *is between-region speciation rates for widespread species AB to speciate into two new species, each in region A and B. e*_*A*_ *and e*_*B*_ *are local extinction rates in region A and B. d*_*AB*_ *and d*_*BA*_ *are dispersal rates from region A into region B and from region B into region A, respectively. For a 2-region GeoSSE model, we have:*

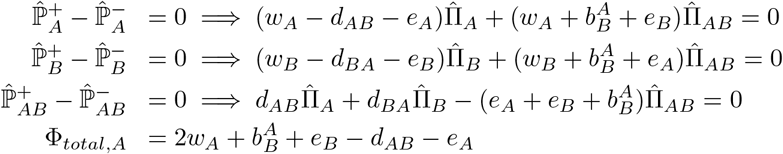

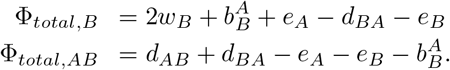

**Proof:** Proof of Corollary S1 is given in Supplement 1.5. □

We demonstrate Corollary S1 in Example S2.

*Example S2* Suppose we have the following stationary frequencies under a 2-region GeoSSE model, 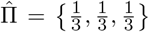. Then, applying Corollary S1, we have two rulesets associated with the frequencies, given as follows:

**Ruleset 1**

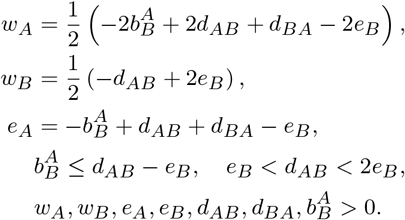

**Ruleset 2**

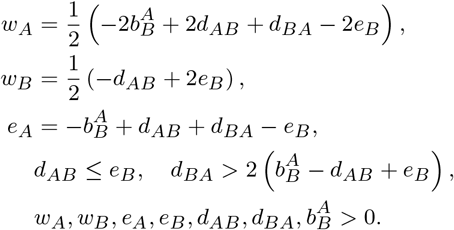

For examples, the following parameters 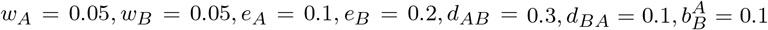 satsify Ruleset 1 but does not satisfy Ruleset 2. On the other hand, 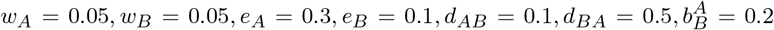 satisfy Ruleset 2 but does not satisfy Ruleset 1. Under both cases, they correspond to the same stationary frequencies 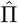 using method given in Soewongsono and Landis (2024).

### 1.5 Proof of Corollary S1

In the context of a 2-region GeoSSE model, we can gain a new species with range state *A* through 1) a within-region speciation event on endemic parent species in state *A* or widespread parent species *AB*, 2) a between-region speciation event on widespread species *AB*, and 3) local extinction event in region *B* by widespread species *AB*. Therefore:

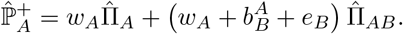

On the other hand, we can lose a new species in *A* through 1) a local extinction event in region *A* and 2) anagenetic change through dispersal event to a widespread species *AB*. Therefore:

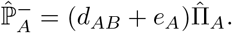

Similarly, we can define:

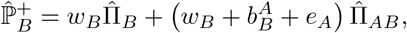

and

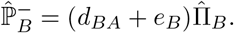

Next, we can gain a widespread species *AB* through a dispersal event from species in region *A* to region *B*, and vice versa. We can lose a widespread species through a local extinction event in region *A* and region *B*, and through a between-region speciation event. Therefore:

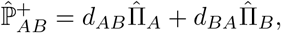

and

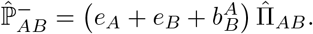

Φ_*total,A*_ and Φ_*total,B*_ represent the flux between all incoming and outgoing rates from events occurring in state *A* and *B*, respectively. In a GeoSSE model, the incoming rates are from within-region speciation on both endemic and widespread species and from between-region speciation and local extinction on widespread species. While, the outgoing rates are from local extinction event on endemic species. Thus:

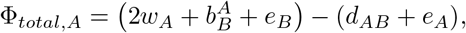

and

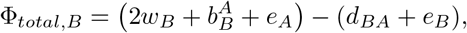

As for the widespread species, we have

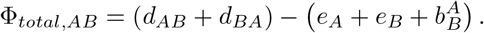

### 1.6 Proof of Lemma 1

The proof follows directly from Theorem 1. Suppose we have an arbitrary pair of symmetric state frequencies namely, 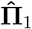 and 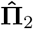. Suppose the rate parameters that correspond to 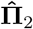 are obtained by re-labelling the indices in rate parameters that correspond to 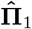. Then, in both patterns, we solve the same system of equations and constraints as described in Theorem 1. Thus, they both return the same solution sets. In other words, both pairs of symmetric state patterns return symmetric rulesets after re-labelling the indices on rate parameters.

### 1.7 Symmetries between rulesets in GeoSSE

*Example S3* Suppose a 2-region GeoSSE model with state space *S* = {*A, B, AB*} with 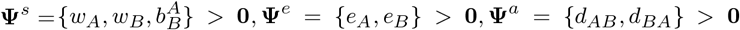. Then, given a pair of symmetric state frequencies as follows:

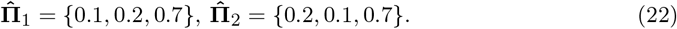

Then, using Corollary S1 (solved using `Mathematica` (Wolfram Research, Inc. 2023)), rulesets that correspond to 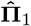 are as follows:

1. **Ruleset 1**

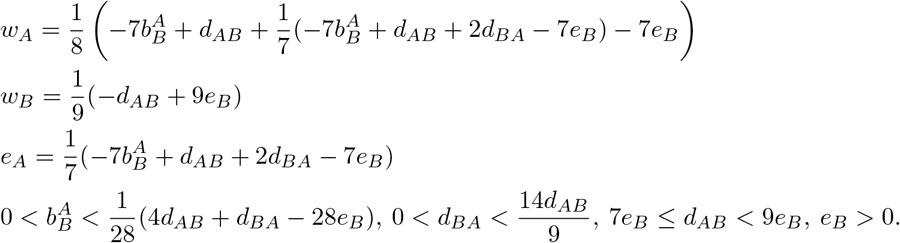
2. **Ruleset 2**

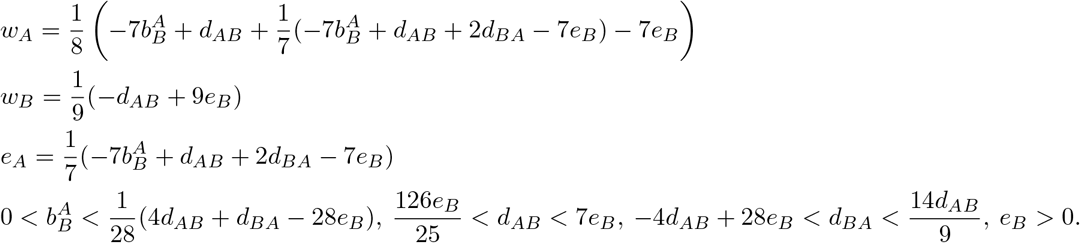

On the other hand, rulesets that correspond to 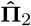 are as follows:

1. **Ruleset 1**

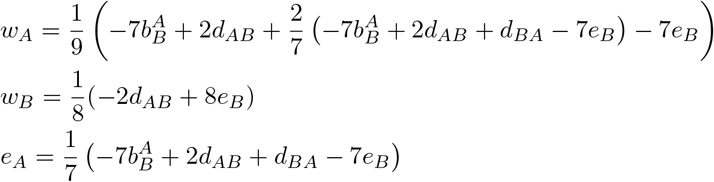

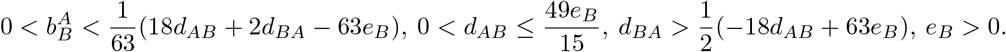
2. **Ruleset 2**

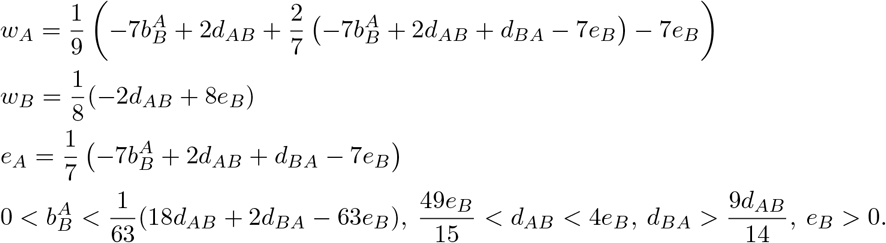

Note that `Mathematica` has a tendency to preserve the order of how expressions are being represented, depending on user’s order of input parameters. This leads to the symmetry between rulesets for a pair of symmetrical stationary state frequencies is being **indirectly** represented. However, the ruleset 1 for 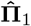 is indeed symmetric to the ruleset 2 for 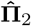. Similarly, ruleset 2 for 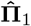 is symmetric to the ruleset 1 for 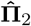. To show the symmetries, we can re-write both rulesets in 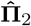 (equivalent to changing the order of input parameters in `Mathematica`) as follows:

1. **Ruleset 1**

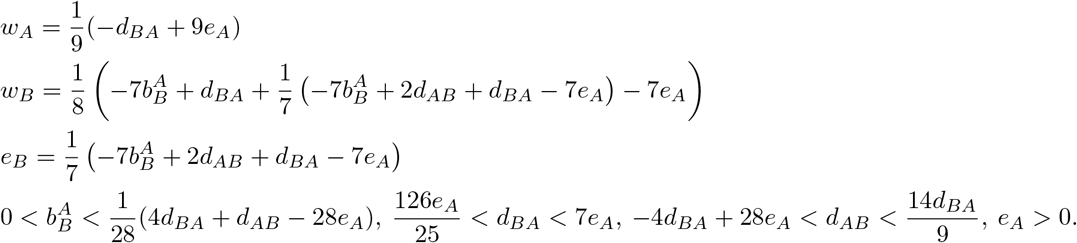
2. **Ruleset 2**

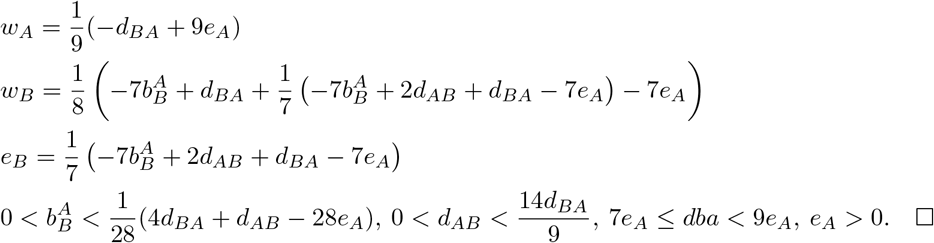

### 1.8 Proof of Corollary 3

From Corollary 1 we have,

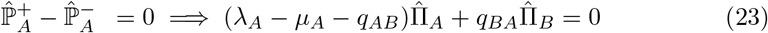

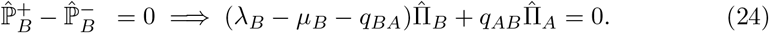

#### Case 1

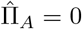

Then, from Eq (23) we have:

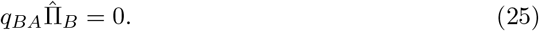

However, since 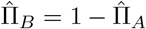, from Eq. (25) we have 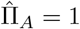 since *q*_*BA*_ ≠ 0. This is a contradiction.

#### Case 2

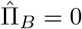

Proof for Case 2 follows directly from proof for Case 1 by re-labelling the rate parameters. By symmetry, 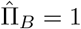 since *q*_*AB*_≠ 0. This is a contradiction. □

### 1.9 Zero ruleset in GeoSSE

#### Corollary S4.

*Suppose we have a 2-region GeoSSE model with state space S* = {*A, B, AB*} *with rate parameters* 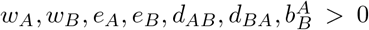. *Given a set of stationary frequencies* 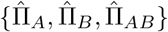, *there is no non-trivial ruleset satisfying the system of equations in Corollary S1 if either*

i. *at least one stationary frequency is zero, or*
ii. 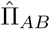 *is equal to exactly one of* 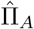 *or* 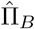.

*Proof* We first show for case (i). From Corollary S1 we have:

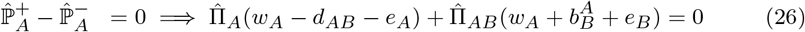

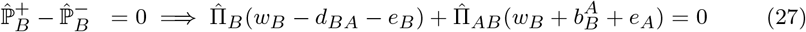

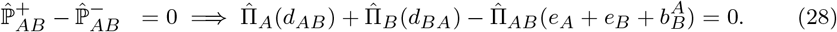

#### Scenario 1

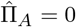

Then, from Eq. (26) we have:

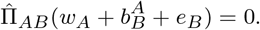

Since 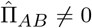 this implies that 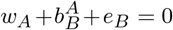. However, by assumption, 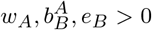.

Thus, it is a contradiction.

#### Scenario 2

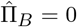

From Eq. (27) we have

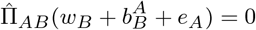

Since 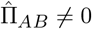, this implies that 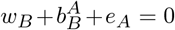. However, by assumption, 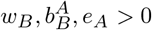.

Thus, it is a contradiction.

#### Scenario 3

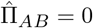

From Eq. (28) we have:

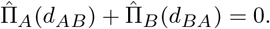

By assumption, *d*_*AB*_, *d*_*BA*_ *>* 0. Thus, 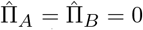. However, this is a contradiction. Next, to show case (ii), we assume that 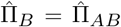, and all stationary frequencies are positive. The proof for 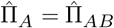 follows by symmetry. From Eq. (27) we have

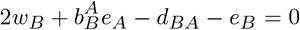

since 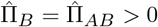. By definition, it implies

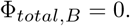

Moreover, Φ_*total,AB*_ = 0 since 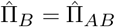. Then, we have

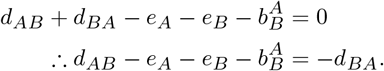

From Eq. (28) we have

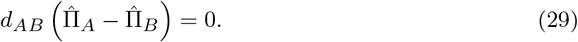

This implies *d*_*AB*_ = 0 since 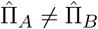. This is a contradiction since all rates are positive. □

### 1.10 Proof of Theorem 2

First, we prove the equalities that define *λ*_*A*_ and *λ*_*B*_ in rulesets 1*a/b* and 2*a/b*. By re-arranging parameters in Eqs. (4) and (5) in Corollary 1, we have:

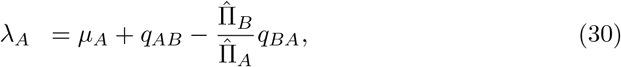

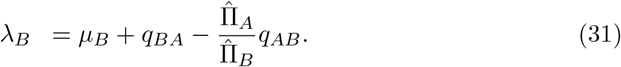

Then, the proof for *λ*_*A*_ and *λ*_*B*_ in rulesets 1*c* and 2*c* follows directly since 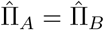.

Next, we consider case 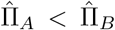 to prove the inequalities in ruleset 1*a*. Let’s suppose we have rate corresponds to species gain in state *A*. That is, *λ*_*A*_ *> µ*_*A*_. Since *λ*_*A*_ *> µ*_*A*_ we have:

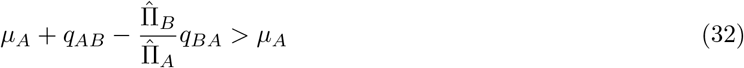

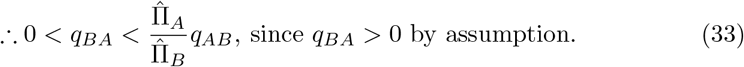

Next, by assumption *λ*_*B*_ *>* 0 we have:

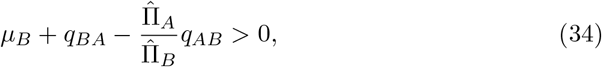

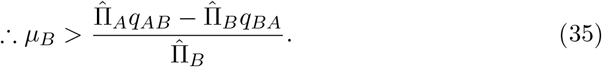

It also follows that *λ*_*B*_ *< µ*_*B*_ by Eqs. (31) and (33). This completes the proof for ruleset 1*a*. Next, assuming we have rate corresponds to species gain in *B*. That is *λ*_*B*_ *> µ*_*B*_. Since *λ*_*B*_ *> µ*_*B*_ we have

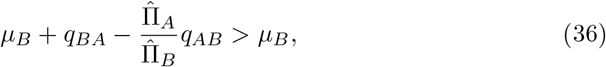

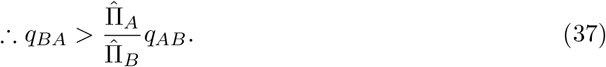

Next, by assumption *λ*_*A*_ we have:

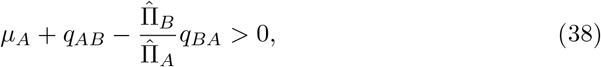

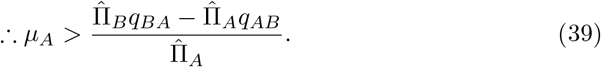

It also follows that *λ*_*A*_ *< µ*_*A*_ by Eqs. (30) and (37). This completes the proof for ruleset 2*a*. Next, for case 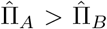, the proof for the inequalities in rulesets 1*b* and 2*b* follow directly by swapping the state indices in the inequalities for rulesets 1*a* and 2*a*, respectively. Next, we consider case 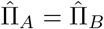 to prove the inequalities in rulesets 1*c* and 2*c*. Consider *λ*_*B*_ *> µ*_*B*_. It follows from Eq. (37) that:

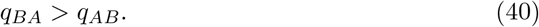

Next, from Eq. (39) we have:

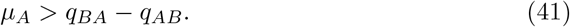

This completes the proof for ruleset 1*c*. Next, assuming *λ*_*A*_ *> µ*_*A*_, it follows from Eq. (33) that:

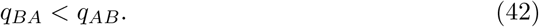

Next, from Eq. (35) we have:

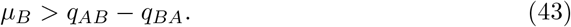

This completes the proof for ruleset 2*c*. □

### 1.11 Proof of Lemma 2

Following the proof in Theorem 2 for ruleset 1*a*, we have *λ*_*A*_ *> µ*_*A*_ and *λ*_*B*_ *< µ*_*B*_. Moreover, we have:

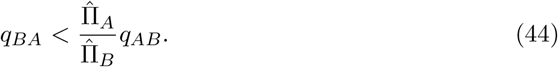

Since ruleset 1*a* corresponds to 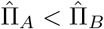, it follows that *q*_*BA*_ *< q*_*AB*_. This completes the proof of the mapping between ruleset 1*a* and scenario 1*a*.

Next, following the proof in Theorem 2 for ruleset 2*a*, we have *λ*_*A*_ *< µ*_*A*_ and *λ*_*B*_ *> µ*_*B*_. Moreover, Eq. (37) still holds regardless of whether *q*_*AB*_ *< q*_*BA*_ or *q*_*AB*_ *> q*_*BA*_. This completes the proof of the mapping between ruleset 2*a* and scenario 2*a*. Next, by symmetry, the proof for scenarios 1*b* and 2*b* follows directly from 1*a* and 2*a*.

Following the proof in Theorem 2 for ruleset 1*c*, we have *λ*_*B*_ *> µ*_*B*_. Since ruleset 1*c* corresponds to 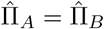, it follows from Eq. (30) that:

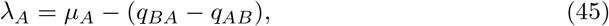

and so, *λ*_*A*_ *< µ*_*A*_ since *q*_*BA*_ *> q*_*AB*_. This completes the proof of the mapping between ruleset 1*c* and scenario 1*c*. Next, following the proof in Theorem 2 for ruleset 2*c*, we have *λ*_*A*_ *> µ*_*A*_. Moreover, since *q*_*BA*_ *< q*_*AB*_ it follows from Eq. (31) that *λ*_*B*_ *< µ*_*B*_. This completes the proof of the mapping between ruleset 2*c* and scenario 2*c*.

Finally, in order to show the other non-trivial scenarios described by the Lemma, it is sufficient to show their existence and their inherent exclusion by the method described in Corollary 1. Let’s assume they are included by the method.

In scenario 3*a* and 4*a*, we have *λ*_*A*_ *> µ*_*A*_ and *λ*_*B*_ *> µ*_*B*_. Therefore, it follows from Eqs. (30) and (31) that:

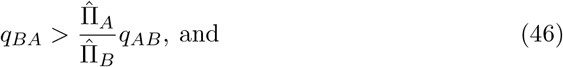

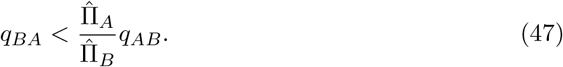

However, this is a contradiction. Hence, using proof-by-contradiction, scenarios 3*a* and 4*a* are excluded by the method and so, we cannot derive explicit rulesets for these scenarios using the method described in Corollary 1. The existence of scenarios 3*a* and 4*a* follows directly from Corollary 2. That is, we have *λ*_*A*_ *> µ*_*A*_ and *λ*_*B*_ *> µ*_*B*_.

Furthermore, 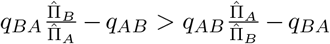 implies 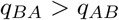, and 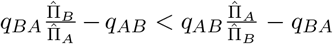 implies *q*_*BA*_ *< q*_*AB*_. This completes the proof for scenarios 3*a* and 4*a*. By symmetry, the proof for scenarios 3*b* and 4*b* also follows in a similar manner.

Next, let’s assume we can derive scenarios 5*a* and 6*a* using method described in Corollary 1. Then, given that *λ*_*A*_ *< µ*_*A*_ and *λ*_*B*_ *< µ*_*B*_, it follows from Eqs. (30) and (31) that:

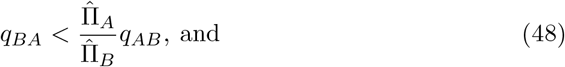

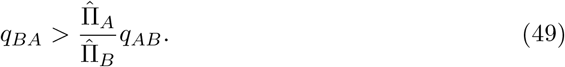

However, this is a contradiction. The existence of scenarios 5*a* and 6*b* follows directly from Corollary 2. That is, we have *λ*_*A*_ *< µ*_*A*_ and *λ*_*B*_ *< µ*_*B*_. Furthermore, 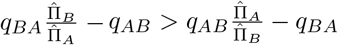 implies 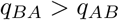, and 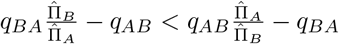 implies *q*_*BA*_ *< q*_*AB*_. This completes the proof for scenarios 5*a* and 6*a*. By symmetry, the proof for scenarios 5*b* and 6*b* also follows in a similar manner.

Next, in Scenario 7*a* we have *λ*_*A*_ − *µ*_*A*_ = *λ*_*B*_ − *µ*_*B*_. Then, using Eqs. (30) and (31) we have:

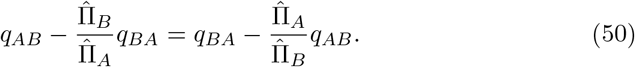

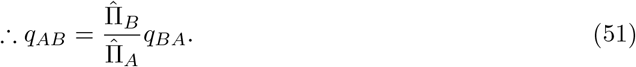

Substitute Eq. (51) to Eq. (4) in Corollary 1 we have:

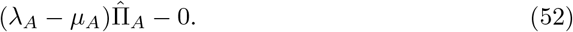

However, since 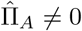. Then, it implies *λ*_*A*_ = *µ*_*A*_, which is a contradiction. However, this scenario does exist. If the rate of species accumulation in both state through speciation are equal, then the direction of the transition rate determines the direction of the stationary class. In this case, since *q*_*AB*_ *> q*_*BA*_, then it follows that 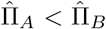. This completes the proof for scenario 7*a*. By symmetry, the proof for scenario 7*b* follows in a similar manner.

For scenario 3*c*, as usual, first we show that there is no ruleset that corresponds to this scenario under the method described in Corollary 1. Then, we show the existence of such scenario. Since *q*_*AB*_ = *q*_*BA*_, it follows from Eq. (4) in Corollary 1 that *λ*_*A*_ = *µ*_*A*_, which is a contradiction. Next, we show the existence of the scenario. By the assumption on the rate parameters under this scenario, it follows that 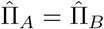 since it is equivalent to a state-independent diversification model. □

### 1.12 Proof of Corollary 4

*Proof* From Corollary 2 we have:

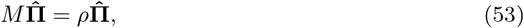

where

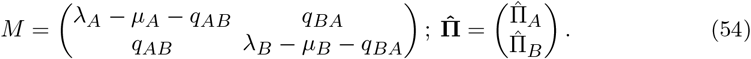

Eq. (53) defines that, at equilibrium, the net contribution to each state is proportional to its abundance and each state grows at the same rate *ρ*. Therefore, the resulting state frequencies are constant over time. Using quadratic formula, the eigenvalues of *M* are given by:

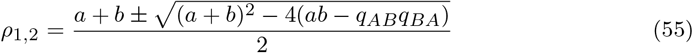

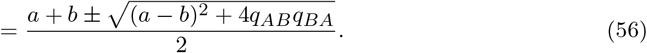

where

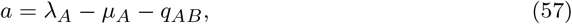

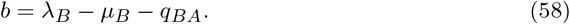

Note that discriminant in Eq. (56) is positive, therefore both eigenvalues are distinct and real. Denote the dominant eigenvalue *ρ* = *ρ*_1_. It follows that the spectral gap is given by:

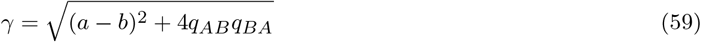

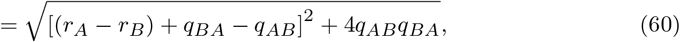

where *r*_*A*_ = *λ*_*A*_ − *µ*_*A*_ and *r*_*B*_ = *λ*_*B*_ − *µ*_*B*_ are net diversification rates. By definition, it follows that the relaxation time is given by:

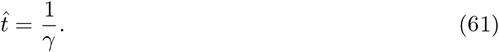

### 1.13 Number of non-trivial scenarios in BiSSE

**Fig. S1:**
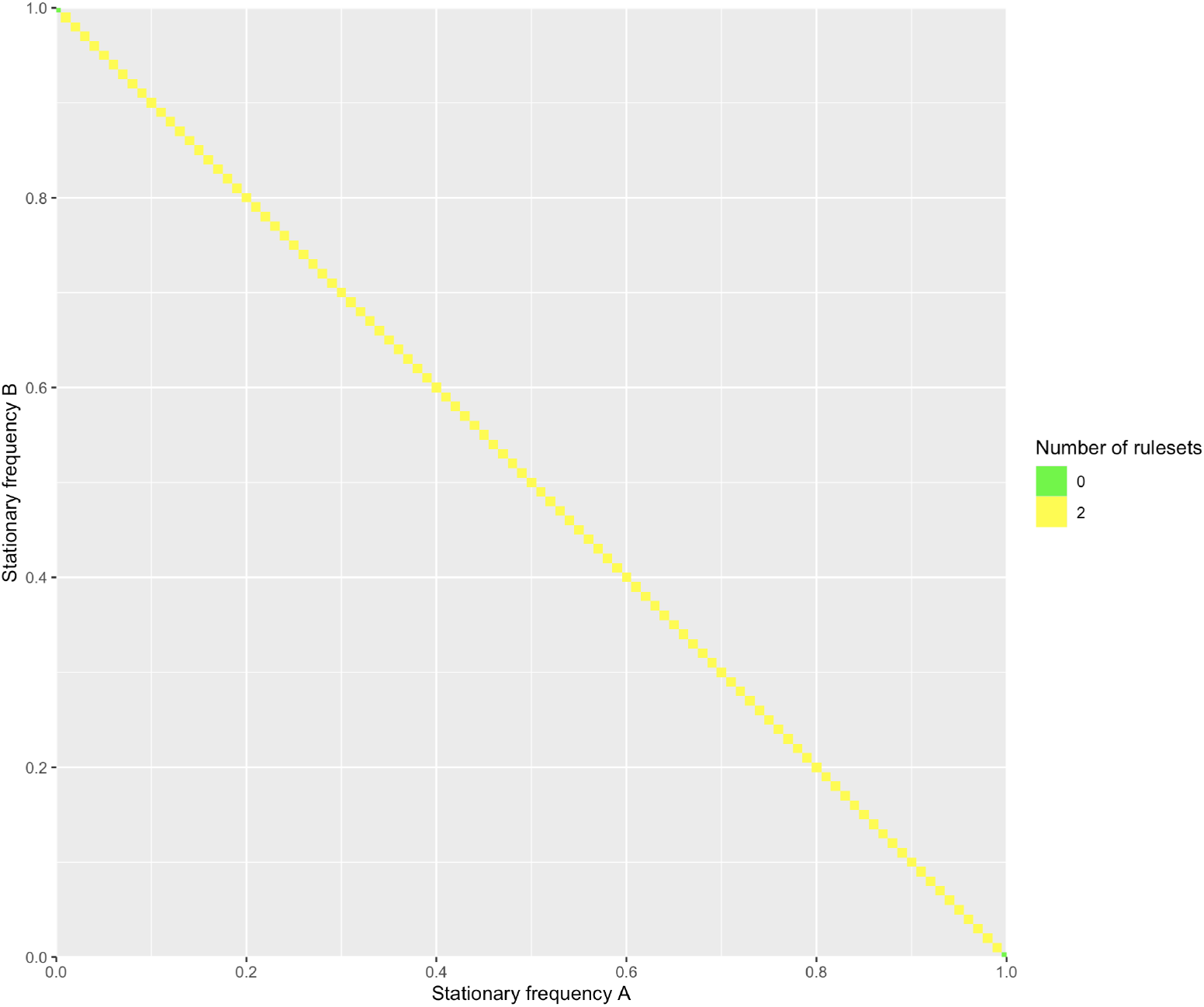
Number of non-trivial rulesets associated with each set of stationary frequencies sampled from a hypercube under the BiSSE model. Because 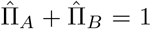, all possible stationary frequency combinations lie along the diagonal. Two non-trivial rulesets exist for 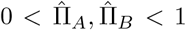, whereas no ruleset exists at the boundary cases 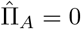 or 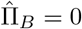 under the assumption that all model rates are positive.

### 1.14 Number of non-trivial scenarios in 3-region GeoSSE

**Fig. S2:**
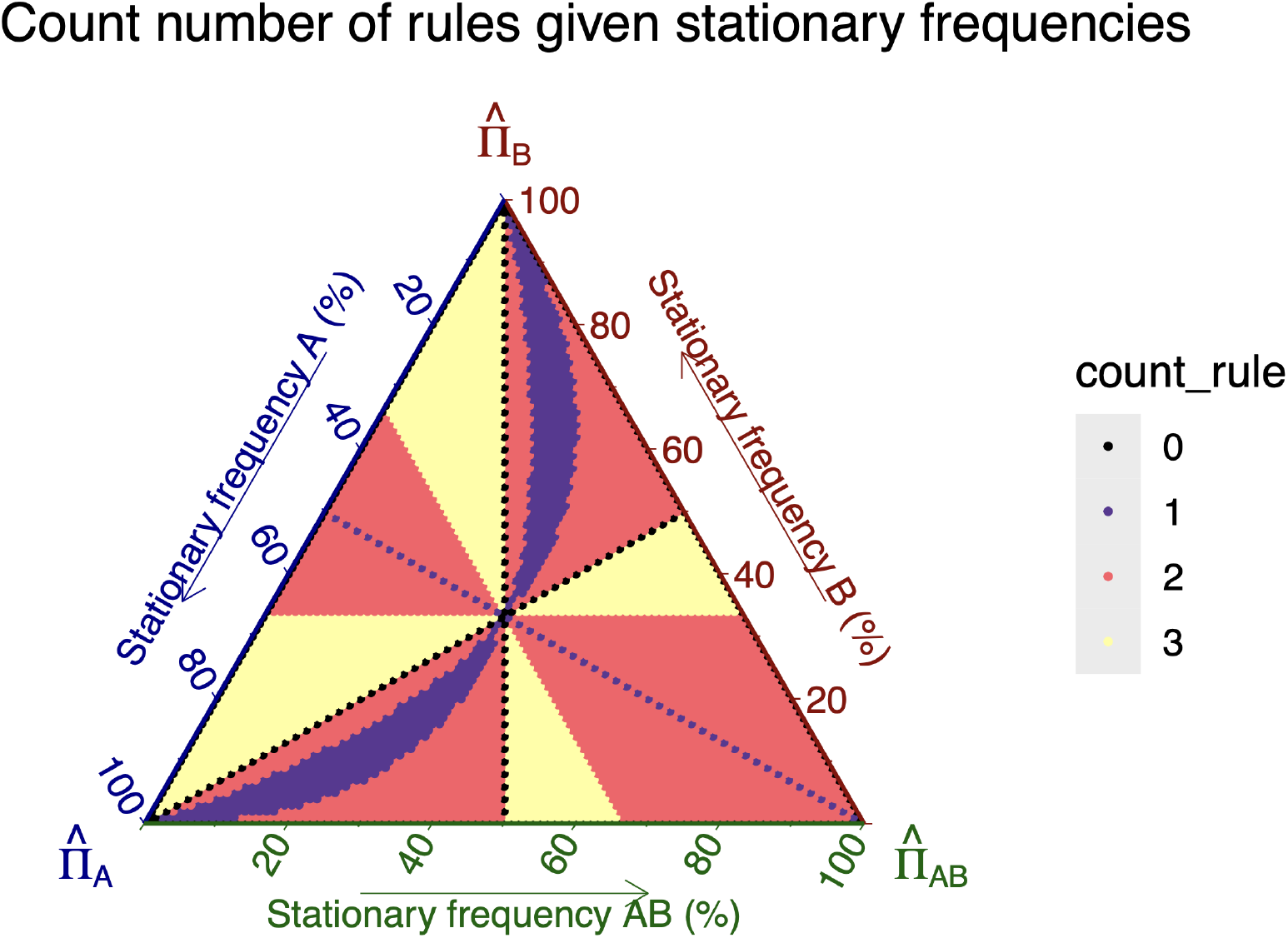
Ternary plot showing the number of non-trivial rulesets associated with each set of stationary frequencies sampled from a hypercube under the 2-region GeoSSE model.

### 1.15 Validation with randomized root states

**Table S1:**
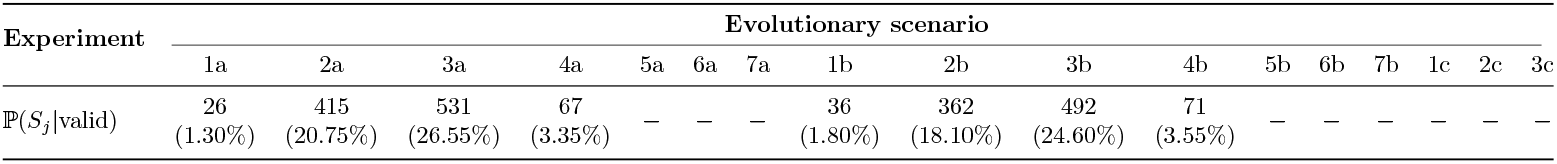
Distribution of true evolutionary scenarios among valid trees, each with 100 extant taxa, under the randomized root validation experiment. The table reports the number of trees, with the relative prevalence in parentheses, for each true scenario among all valid trees. The initial state was randomly assigned to state *A* or *B* with equal probability. Trees were simulated under the complete-noreject setting without rejection based on tip-state frequencies.

### 1.16 BiSSE simulation experiments

**Table S2:**
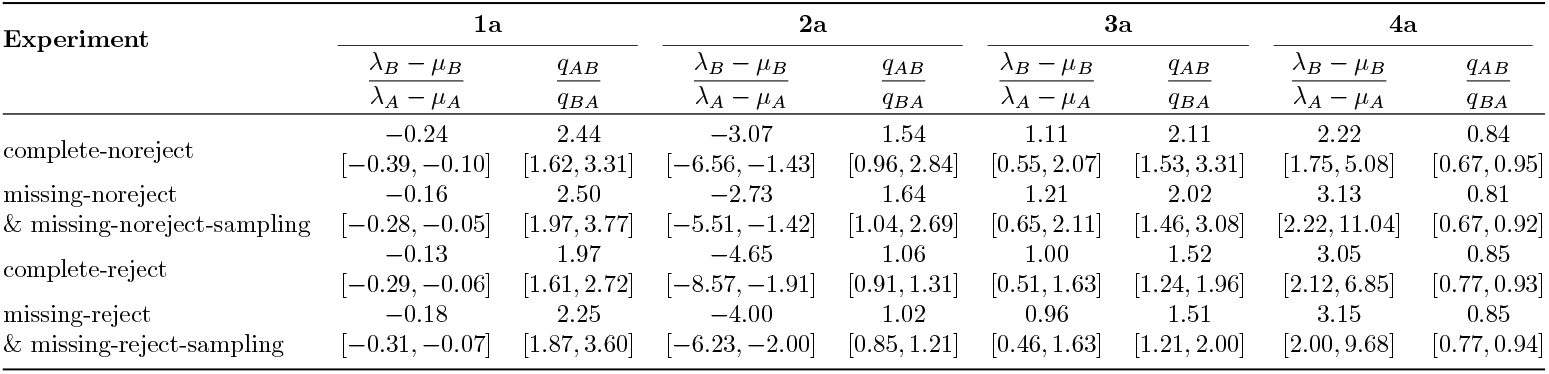
Distribution of the true parameter relationships for simulated trees classified into evolutionary scenarios 1a–4a. The single value reports the median, whereas the interval reports the first and third quartiles of each parameter ratio.

**Table S3:**
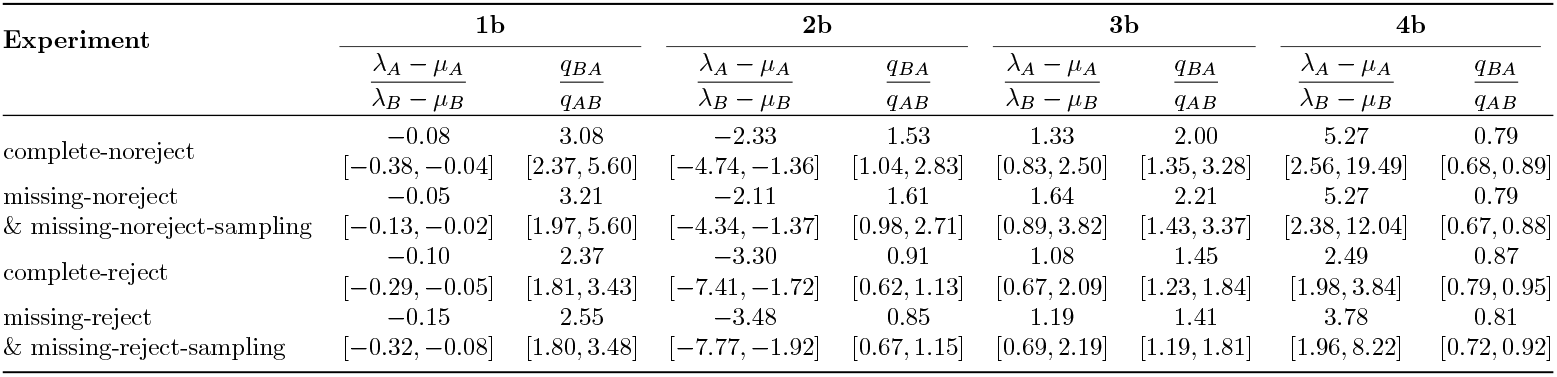
Distribution of the true parameter relationships for simulated trees classified into evolutionary scenarios 1b–4b. The single value reports the median, whereas the interval reports the first and third quartiles of each parameter ratio.

### 1.17 Spectral gap analysis on simulated dataset

**Fig. S3:**
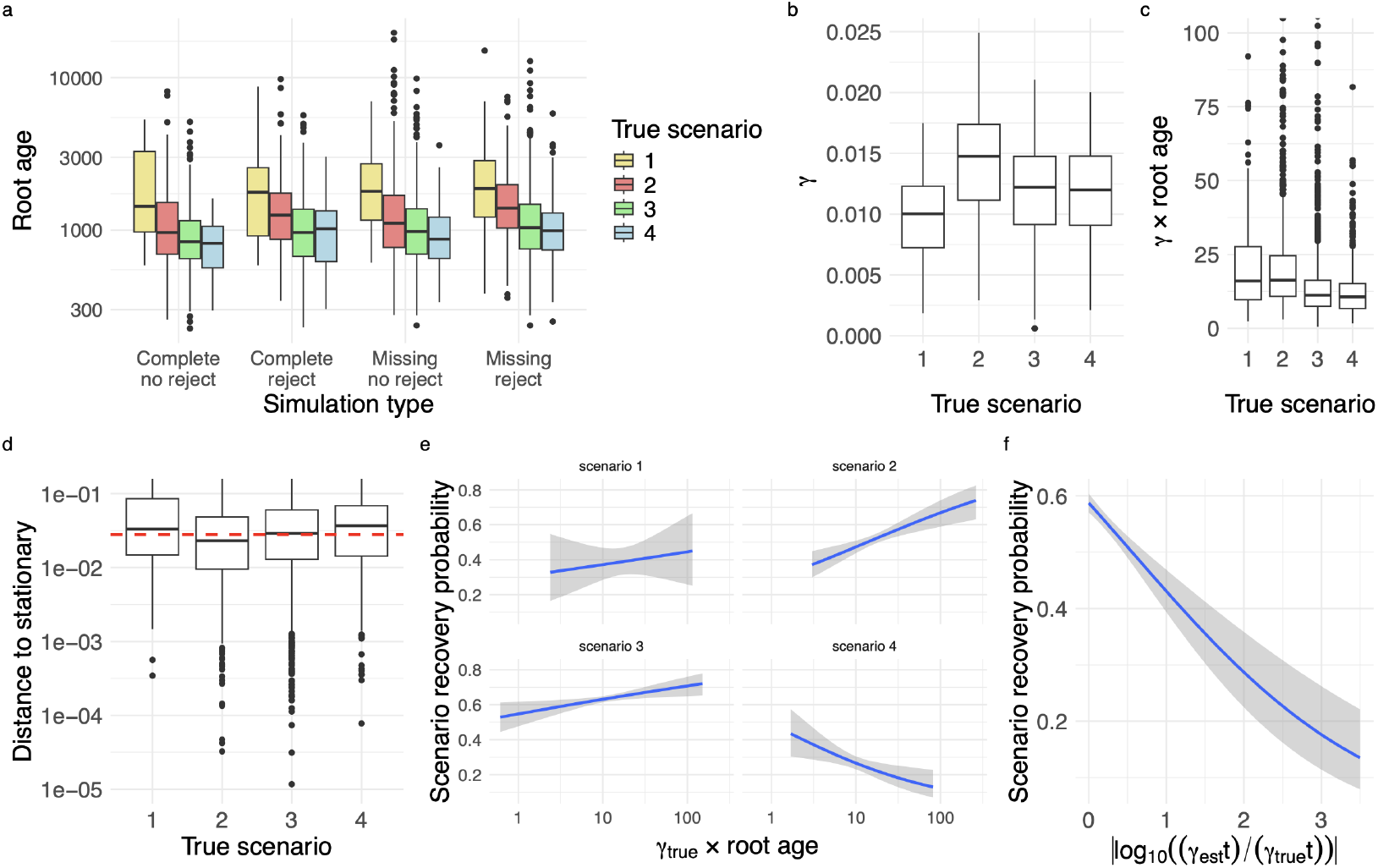
Convergence properties and evolutionary-scenario recovery for datasets simulated using slow parameter rates. (a) Distribution of root ages across the four simulation settings, as described in Table 1, grouped by true evolutionary scenario. (b) Distribution of the spectral gap, *γ*, computed using the true generating parameters for each evolutionary scenario. (c) Distribution of the product *γt*, where *t* denotes the root age. (d) Distance between the observed tip state frequencies and the stationary frequencies under the true generating process for each evolutionary scenario. (e) Probability of correctly recovering the true evolutionary scenario as a function of *γ*_true_*t*. Corresponding “a” and “b” directionality classes are pooled by scenario number for fitting and visualization, but recovery requires an exact match between the estimated and true evolutionary scenario, including the directionality class (e.g., 2a is correctly recovered only when inferred as 2a, not 2b). (f) Probability of correctly recovering the true evolutionary scenario as a function of the absolute log-ratio between the estimated and true spectral gaps, |log_10_(*γ*_est_*/γ*_true_) |. Lines in panels (e) and (f) show fitted binomial logistic regressions, with shaded regions indicating 95% confidence intervals. For all boxplots, center black lines indicate medians, boxes span the interquartile range (IQR), whiskers extend to the most extreme observations within 1.5*×* the IQR, and points beyond the whiskers indicate outliers.

### 1.18 Confusion matrix using trees with missing taxa and without rejection on imbalanced tip states

**Table S4:**
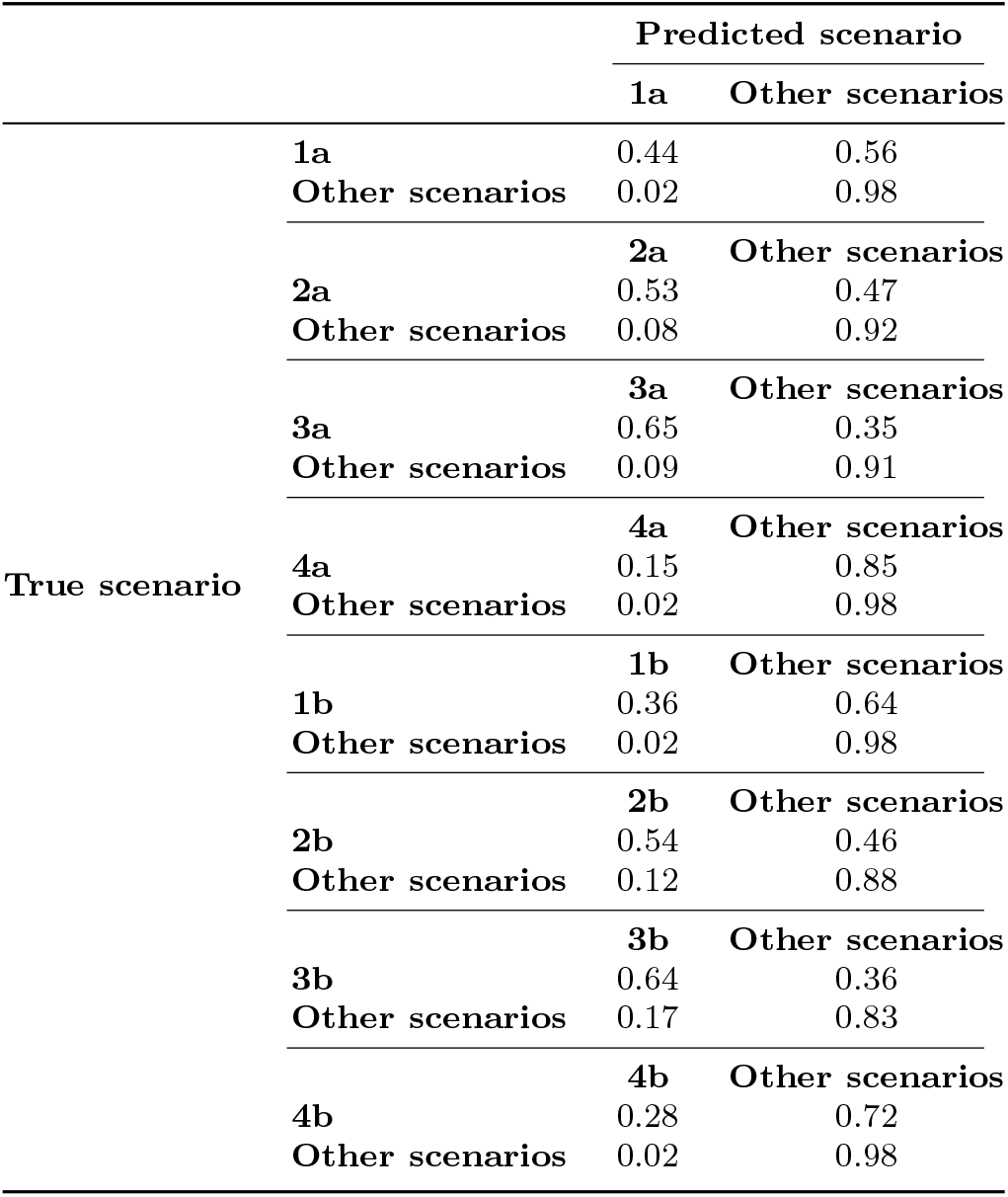
Prediction accuracy for each true scenario versus all other scenarios using the missing-noreject experiment. Scenarios 5a/b, 6a/b, 7a/b, 1c, 2c, and 3c are omitted because they do not have valid tree representation

### 1.19 Confusion matrix using trees with missing taxa and without rejection on imbalanced tip states and account for sampling proportion

**Table S5:**
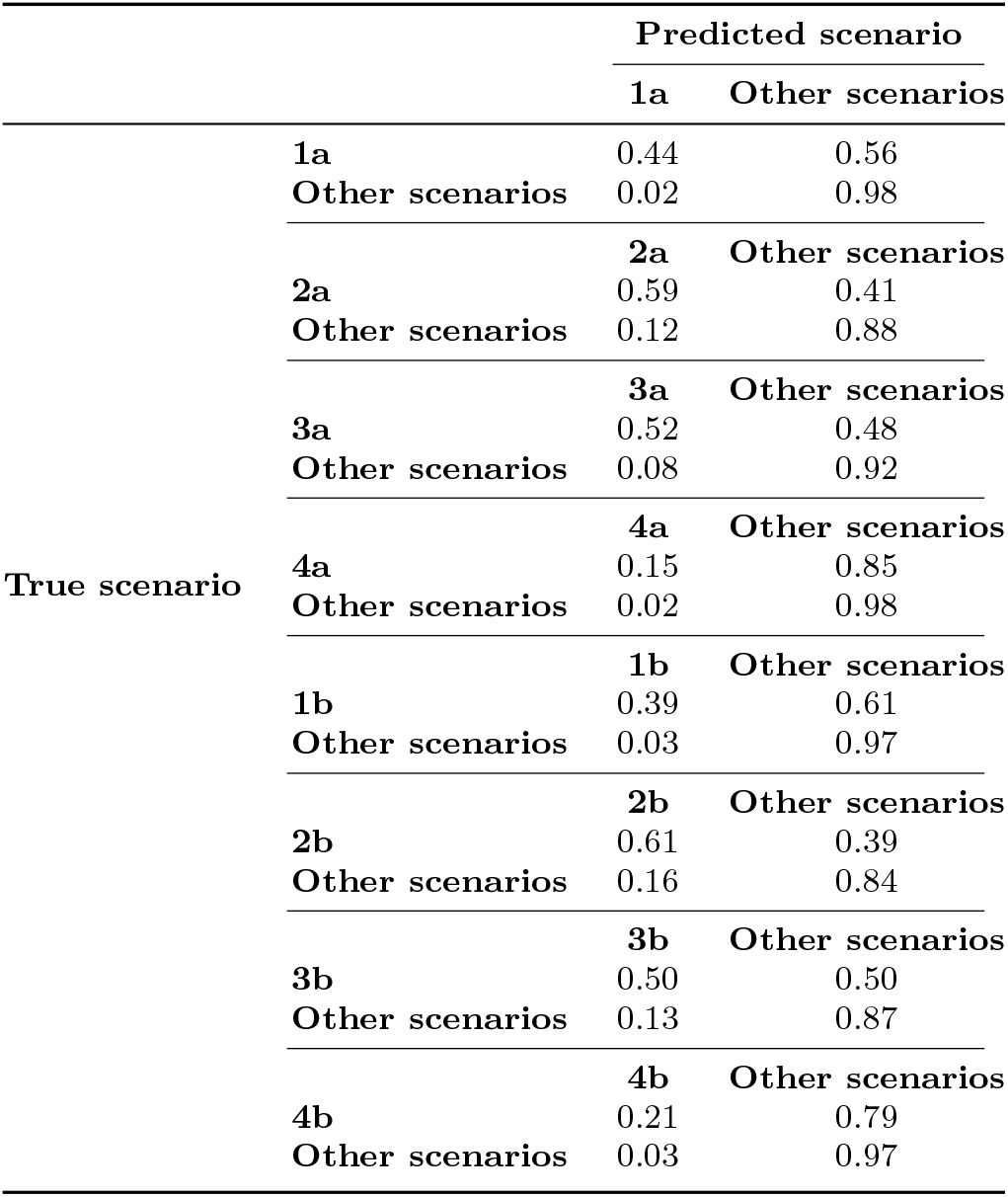
Prediction accuracy for each true scenario versus all other scenarios using the missing-noreject-sampling experiment. Scenarios 5a/b, 6a/b, 7a/b, 1c, 2c, and 3c are omitted because they do not have valid tree representation.

### 1.20 Confusion matrix using trees with complete sampling and with rejection on imbalanced tip states

**Table S6:** Prediction accuracy for each true scenario versus all other scenarios using the complete-reject experiment. Scenarios 5a/b, 6a/b, 7a/b, 1c, 2c, and 3c are omitted because they do not have valid tree representation.

|  |  | Predicted scenario |  |
| --- | --- | --- | --- |
|  |  | 1a | Other scenarios |
| True scenario | 1a | 0.45 | 0.55 |
|  | Other scenarios | 0.06 | 0.94 |
|  |  | 2a | Other scenarios |
|  | 2a | 0.51 | 0.49 |
|  | Other scenarios | 0.10 | 0.90 |
|  |  | 3a | Other scenarios |
|  | 3a | 0.50 | 0.50 |
|  | Other scenarios | 0.10 | 0.90 |
|  |  | 4a | Other scenarios |
|  | 4a | 0.29 | 0.71 |
|  | Other scenarios | 0.03 | 0.97 |
|  |  | 1b | Other scenarios |
|  | 1b | 0.42 | 0.58 |
|  | Other scenarios | 0.06 | 0.94 |
|  |  | 2b | Other scenarios |
|  | 2b | 0.55 | 0.45 |
|  | Other scenarios | 0.13 | 0.87 |
|  |  | 3b | Other scenarios |
|  | 3b | 0.46 | 0.54 |
|  | Other scenarios | 0.11 | 0.89 |
|  |  | 4b | Other scenarios |
|  | 4b | 0.34 | 0.66 |
|  | Other scenarios | 0.04 | 0.96 |

### 1.21 Confusion matrix using trees with missing taxa and with rejection on imbalanced tip states

**Table S7:**
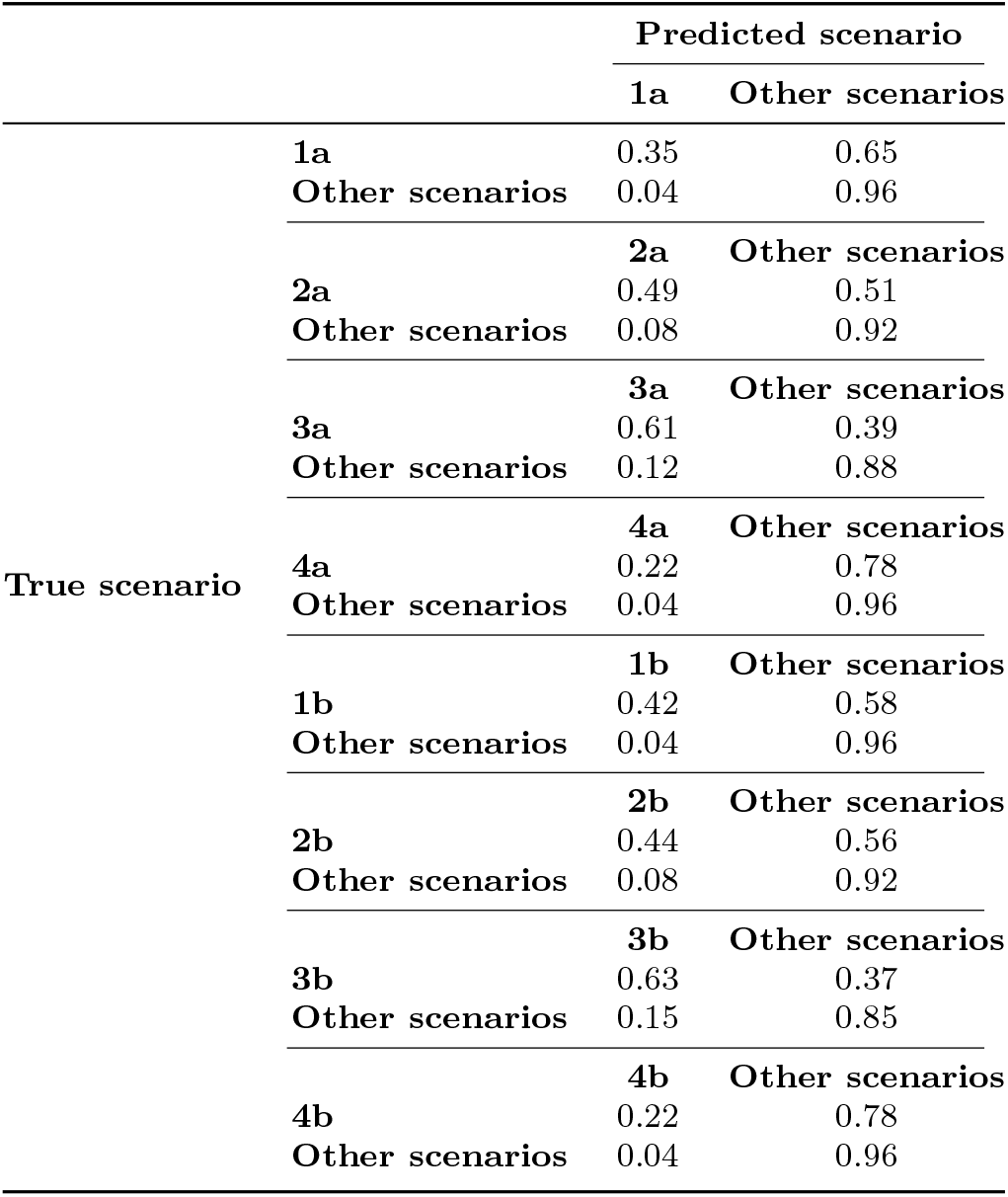
Prediction accuracy for each true scenario versus all other scenarios using the missing-reject experiment. Scenarios 5a/b, 6a/b, 7a/b, 1c, 2c, and 3c are omitted because they do not have valid tree representation.

### 1.22 Confusion matrix using trees with missing taxa and with rejection on imbalanced tip states and account for sampling proportion

**Table S8:**
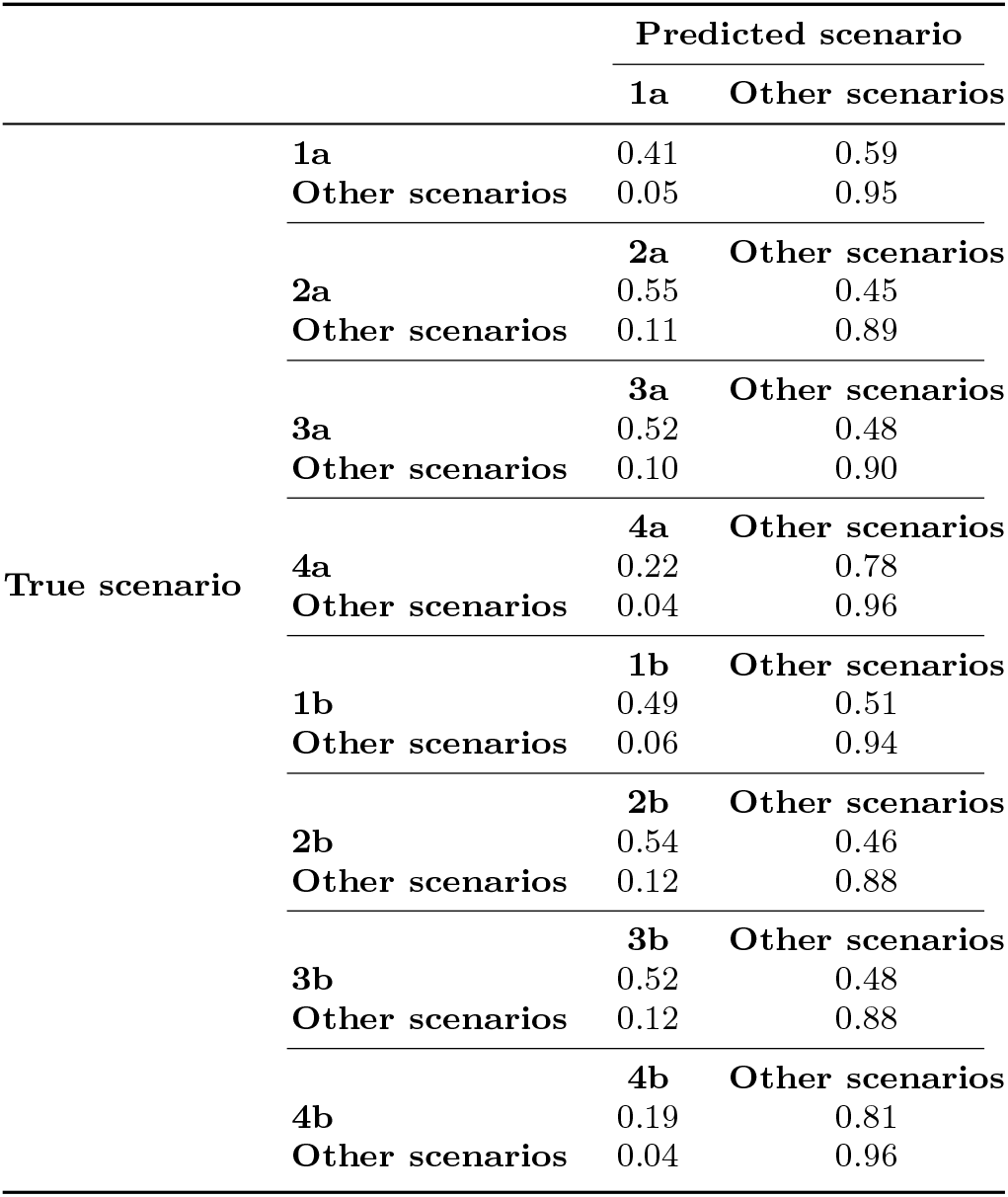
Prediction accuracy for each true scenario versus all other scenarios using the missing-reject-sampling experiment. Scenarios 5a/b, 6a/b, 7a/b, 1c, 2c, and 3c are omitted because they do not have valid tree representation.

